# Filopodia-based contact stimulated collective migration drives tissue morphogenesis

**DOI:** 10.1101/2020.10.19.345082

**Authors:** Maik C. Bischoff, Sebastian Lieb, Renate Renkawitz-Pohl, Sven Bogdan

## Abstract

Cells migrate collectively to form tissues and organs during morphogenesis. Contact inhibition of locomotion (CIL) drives collective migration by inhibiting lamellipodial protrusions at cell-cell contacts and promoting polarization at the leading edge. Here, we report on a CIL-related collective cell behavior of myotubes that lack lamellipodial protrusions, but instead use filopodia to move as a cohesive cluster in a formin-dependent manner. Genetic, pharmacological and mechanical perturbation analyses reveal essential roles of Rac2, Cdc42 and Rho1 in myotube migration. They differentially control not only protrusion dynamics but also cell-matrix adhesion formation. Here, active Rho1 GTPase localizes at retracting free edge filopodia. Rok-dependent actomyosin contractility does not mediate a contraction of protrusions at cell-cell contacts but likely plays an important role in the constriction of supracellular actin cables.

We propose that contact-dependent asymmetry of cell-matrix adhesion drives directional movement, whereas contractile actin cables contribute to the integrity of the migrating cell cluster.

## Introduction

The ability of cells to migrate as a collective is crucial during tissue morphogenesis and remodeling^1,2^. The molecular principles of collective cell migration share features with the directed migration of individual cells. The major driving forces in migrating single cells are Rac-mediated protrusions of lamellipodia at the leading edge, formed by Arp2/3 complex dependent actin filament branching and Rho-dependent actomyosin driven contraction at the cell rear^3,4^. Cells can migrate directionally in response to a variety of chemical cues, recognized by cell surface receptors that initiate downstream signaling cascades controlling the activity or recruitment of Rho GTPases. Directional cell locomotion is also controlled by mechanical stimuli such as upon cell-cell contact^5–7^. A well-known phenomenon is contact inhibition of locomotion (CIL), whereby two colliding cells change direction after coming into contact ^8,9^. Mayor and colleagues provided first mechanistic evidence how CIL might act *in vivo* as the driving force to polarize neural crest cells that derived from the margin of the neural tube and disperse by migration during embryogenesis^10,11^.

In neural crest cells, CIL involves distinct stages of cell behavior including cell-cell contact, protrusion inhibition, repolarization, contraction and migration away from the collision^12^. The initial cell-cell contact requires the formation of transient cadherin-mediated cell junctions. Once the cells come in close contact, a disassembly of cell-matrix adhesion near the cell-cell contact and the generation of new cell-matrix adhesions at the free edge occur. Such mechanical crosstalk between N-cadherin-mediated cell-cell adhesions and integrin-dependent cell-matrix adhesions has been recently described *in vivo* during neural crest cell migration in both *Xenopus* and zebrafish embryos^13^. However, the loss of cell-matrix adhesions at cell contacts alone is not sufficient to drive CIL. A subsequent repolarization of the cells away from the cell-cell contact and thereby the generation of new cell-matrix adhesions and protrusions at the free edge are required to induce cell migration away from the collision. In neural crest cells, this depends on the polarized activity of the two Rho GTPases, Rac1 and RhoA^14^. A model of CIL has been proposed in which a contact-dependent intracellular Rac1/RhoA gradient is formed that generates an asymmetric force driving directed cell migration^15^. N-cadherin binding triggers a local increase of RhoA and inhibits Rac1 activity at the site of contact^14,16^. Thus, Rac1-dependent protrusions become biased to the opposite end of the cell-cell contact and cells migrating away from the collision.

Overall, CIL has been successfully used to explain contact-dependent collective migration of loose clusters of mesenchymal cells such as neural crest cells and hemocytes^12^, but it is still unclear whether mechanisms governing CIL might also contribute to the migratory behavior of cohesive cell clusters or epithelia^5,7^.

Here, using an integrated live-cell imaging and genetic approach, we identified a CIL related, contact-dependent migratory behavior of highly cohesive nascent myotubes of the *Drosophila* testis. Myotubes lack lamellipodial cell protrusions, but instead form numerous large filopodia generated at both N-cadherin-enriched cellular junctions at cell-cell contacts and integrin-dependent cell-matrix sites at their free edge. Filopodia-based myotube migration requires formins and the Rho family small GTPases Rac2, Cdc42 and RhoA, whereas the Arp2/3 complex and its activator, the WAVE regulatory complex (WRC) seem only to contribute to filopodia branching. Rac2 and Cdc42 differentially control not only protrusion dynamics but also cell-matrix adhesion formation. Unlike CIL, RhoA is not activated at cell-cell contacts, but rather gets locally activated along retracting protrusions. Genetic and pharmacological perturbation analysis further revealed an important requirement of Rho/Rok-driven actomyosin contractility in myotube migration.

In summary, we propose a model in which N-cadherin-mediated contact dependent asymmetry of cell-matrix adhesion acts as a major switch to drive cell movement towards the free space, whereas contractile actin cables contribute to the integrity of the migrating cell cluster.

## Results

### Long-term live imaging of *Drosophila* smooth-like testes muscles as a new collective cell migration model

At 24h after puparium formation (APF), both testes lay free in the body cavity (Figure 1a). The genital disc provides the myoblasts and other somatic parts of the reproductive system such as the seminal vesicles^17,18^. Testes myoblasts adhere to the epithelium of the seminal vesicles (Figure 1a, sv) and fuse to small syncytia shortly before the connection between seminal vesicles and terminal epithelia (Figure 1a, te) has been formed (Figure 1a, b^19,20^). Between 28-30h APF this connection has been established (Figure 1, see arrow between a and b). At 30 h APF nascent myotubes (Figure 1b, mt in red) start to migrate beneath the pigment cell layer (Figure 1b, pc) to and along the testes towards the apical end (Figure 1b ^21^). At 40 h APF, myotubes cover the whole pupal testis as a thin muscular sheet^22^.

**Figure 1.**
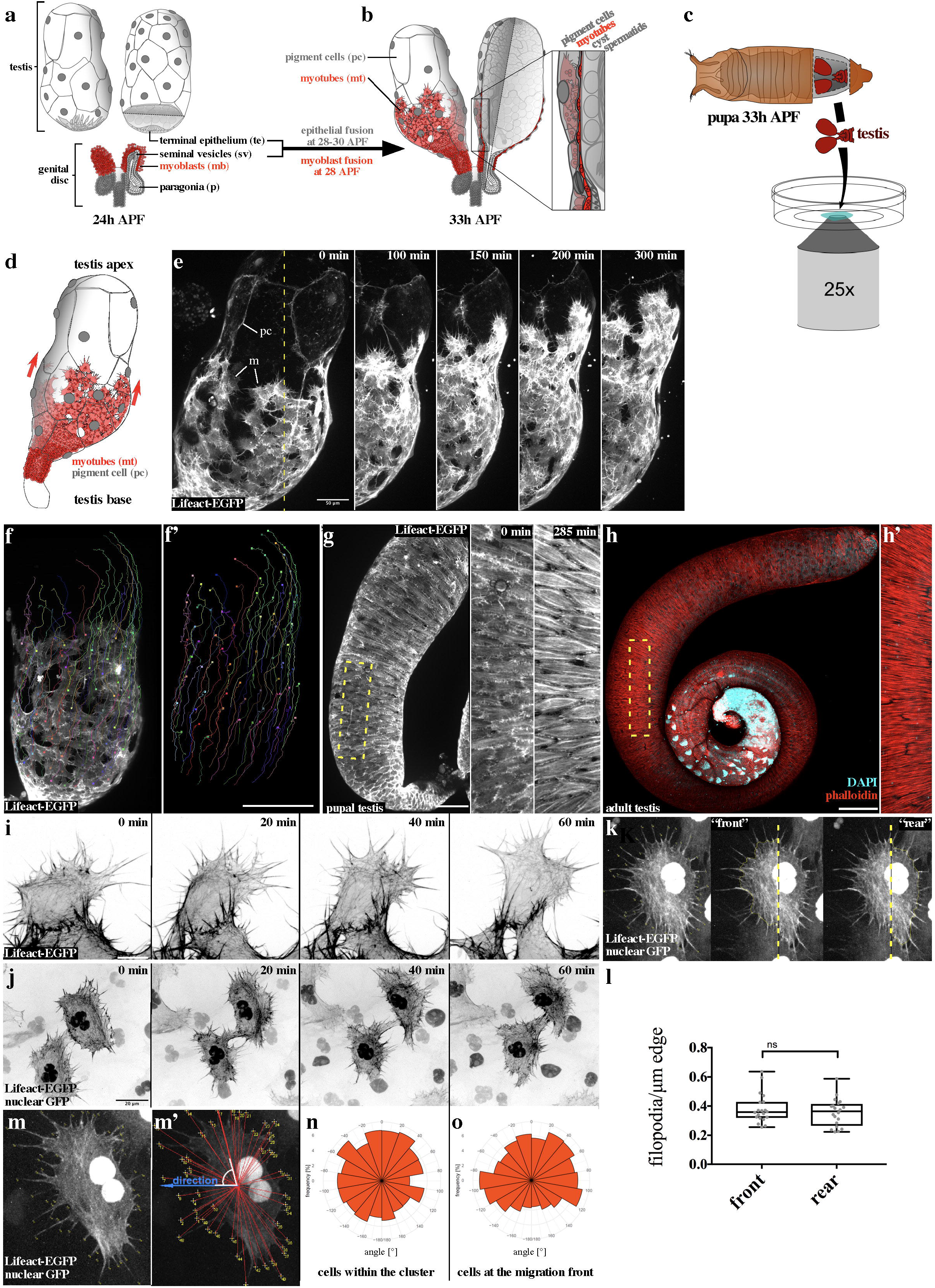
Myotubes form numerous filopodial membrane protrusions instead of lamellipodia and migrate collectively onto the testis. **a, b.** Graphics of the *Drosophila* testis at 24h and 33 h after pupae formation (APF). **a.** Myoblasts (mb, red) arising from the genital disc, adhere to the epithelium of the seminal vesicles (sv) and fuse to small syncytia shortly before the connection between seminal vesicles and terminal epithelia (te). **b.** After epithelial fusion nascent myotubes (mt) migrate between the basal lamina separating the testicular cyst cells and a layer of pigment cells (pc) from the testis base towards the apex. **c.** Schematic of the *ex vivo* technique enabling life imaging of *Drosophila* testis development with a spinning disc microscope. **d.** Only one testis of the pair (compare to c) which was prepared is depicted, as in 25x magnification (compare to d) only one testis can be seen. **e.** Wild type testis 33 h APF 350 min in *ex vivo* culture. UAS-LifeAct-EGFP was driven using the *htl-Gal4* driver line, which promotes expression in migrating myotubes (m) and pigment cells (pc). The dashed line in “0 min” represents the area depicted in 100–420 min. Scale bar, 50 μm. **f, f’.** Migration of myotubes was tracked using the Imaris software. *mef2-Gal4* was used to drive UAS-LifeAct-EGFP, expressed only in myotubes. **f.** An overlay of microscopic data and track data are shown. Source data are provided as a Source Data file. **f’.** Only track data is shown. Scale bar, 100 μm. **e.** Top view of a testis 46 h APF in 420 min *ex vivo* culture. *mef2-Gal4* drives UAS-LifeAct-EGFP expression. **g.** Subsequent to migration, testis myotubes start to encircle the testis, generating ring muscles. The testis starts to change its shape. The dashed line in h represents the area depicted in 0/285 min. Scale bar, 100 μm. **h.** Confocal image of an adult testis stained with phalloidin and DAPI. Due to constriction by building muscles in pupal development, the testis gained its typical coiled shape ^30^ Scale bar, 100 μm; close-up in **h’**. **i.** Close-up of myotubes at the front edge of the migrating sheet 60 min in *ex vivo* culture. *mef2-Gal4* drives UAS-LifeAct-EGFP expression. Cells at the front of the migrating cluster appear like the cells within the cluster depicted in h, as they project filopodia-like structures in all directions. The actin cytoskeleton appears in stress fiber-like thick bundles. Scale bar 10 μm. **j.** Close-up of two myotubes during migration 60 min in *ex vivo* culture. *beatVC-Gal4* promotes expression of UAS-LifeAct-EGFP and UAS-GFP-nls in a mosaic fashion, allowing for the analysis of single cells within the migrating sheet. Nuclei of neighboring cells are marked by yellow asterisks. Even cells within the cluster, enwrapped by neighboring cells, appear to have filopodia-like protrusions and a general mesenchymal phenotype. Scale bar, 20 μm. **l.** Quantification of filopodia number per cell edge length in cells at the migration front. Source data are provided as a Source Data file. Directionality of filopodia of cells was quantified by measuring the orientation angle as illustrated in **m** and **m’**. Quantification revealed no strong bias in filopodia direction neither of cells **n.** within the myotube cluster nor **o.** of cells at the migration front. Source data are provided as a Source Data file.

To better understand how myotubes cover the testis, we established a protocol for *ex vivo* organ cultivation and long-term imaging (7 h) of isolated 33h APF pupal testes (Figure 1c). We used the muscle-specific *mef2*-Gal4 or the *heartless*-Gal4 (*htl*-Gal4) driver to express a UAS-LifeAct-EGFP transgene either in myotubes or in both, myotubes and pigment cells respectively (see supplementary figure S2a). This method provides an excellent experimental system for studying the highly dynamic migratory cell behavior of myotubes and to visualize their actin-rich protrusions over several hours at high resolution. Spinning disc live imaging microscopy of 33 h APF old testes onwards revealed that myotubes migrate collectively on an ellipsoid surface constrained by the outermost layer of pigment cells and the basal membrane enclosing the inner cysts (Figure 1d, e; supplementary movie M1). To better track the migratory behavior of individual cells within the cell cluster we additionally labeled the cells by co-expression of the membrane marker mCD8-RFP enabling precise 4D (xyz and t) trajectory mapping using the Imaris software (Figure 1f; supplementary movie M2). Since all mathematical directionality descriptors for 2D migration (biased angle, persistence angle, straightness, etc.) are based on Euclidean geometry, we had to transform our 3D(+time) datasets into corresponding 2D(+time) datasets for precise cell quantification. A simple xy-projection would neglect curvature and lead to wrong results. Preexisting tools using unwrapping algorithms and Riemannian manifold learning were not compatible with our system^23^. Instead of an unwrapping algorithm fit for every kind of surface, but with some restrictions in angle and distance accuracy, we developed a Mercator-projection based process, which allows for high angle-accuracy but neglects distances (illustrated in supplementary figure S3 b-f).

Dissecting the cell trajectories of wild type myotubes revealed a directional cell behavior with maximal cell movement into the base-apex direction with a speed about 0.37 μm/min over a distance of about 130 μm (Figure 1f’, Figure 4q, supplementary figure S2g; supplementary movie M2). Once myotubes reached the testis apex they started to elongate and form large actin bundles that aligned perpendicular to the pupal testis surface (Figure 1g, supplementary movie M1, middle). After completing pupal development, myotubes form a densely packed muscle sheath surrounding the elongated, tubular adult testis (Figure 1h, h’).

### Myotube migration depends on formin-dependent filopodial membrane protrusions

Strikingly, migrating myotubes largely lacked lamellipodial protrusions, but instead formed numerous filopodia-like protrusions (from here on referred to as filopodia; Figure 1i; supplementary movie M3). Expression of LifeAct-EGFP together with a nuclear targeted EGFP transgene in myotubes in a mosaic-like fashion further showed that myotubes also formed prominent filopodial protrusion between neighboring cells (Figure 1j.; supplementary movie M4). To better characterize the distribution of filopodia in these cells, we quantified the directionality of filopodia of cells by measuring the orientation angle as illustrated in figure 1m and m’. This analysis revealed no strong bias in filopodia generation or directionality in cells within the cluster (Figure 1n) and surprisingly also at the front edge of the cluster (Figure 1o). To statistically analyze this, we differentiate the filopodia (in cells at the front edge) in those which are assembled at the cell front (pointing to the testis apex) and those at the “rear” (pointing to the testis base; Figure 1kl). To account for irregular cell shapes, we calculated the density (number/μm) by measuring edge length. There was no significant difference in filopodia density between front and rear. Thus, the directionality of collective myotube migration cannot be simply predicted by filopodia number or direction.

We next determined how central actin nucleators such as formins and the Arp2/3 complex contribute to filopodia formation and myotube migration. Treatment with the specific Arp2/3 inhibitor CK-666^24^ did not strongly affect the overall cell cluster morphology compared to control cells incubated with DMSO (Figure 2a, b and c; supplementary movie M5). Likewise, cells depleted of the *arp3* subunit or *wave* by RNA interference (RNAi) showed moderate changes in cell morphology despite prominent fusion defects (see mononucleated myotubes marked by co-expression of the mCD8-RFP marker excluded from the nuclei in figure 2e; supplementary movie M6). Similar to CK666 treatment, *arp3* and *wave* depleted cells were still able to migrate persistently in a directed fashion (Figure 2b-e’; supplementary figure S2h. However, cells depleted of the *arp3* subunit or treated with CK666 showed a significantly reduced migration speed and distance along the x-axis (compare migratory tracks in figure 2a, b and d; quantification in figure 2h supplementary figure S2g). Thus, the Arp2/3-WRC pathway promotes motility, but seems to be dispensable for directed migration of myotubes.

**Figure 2.**
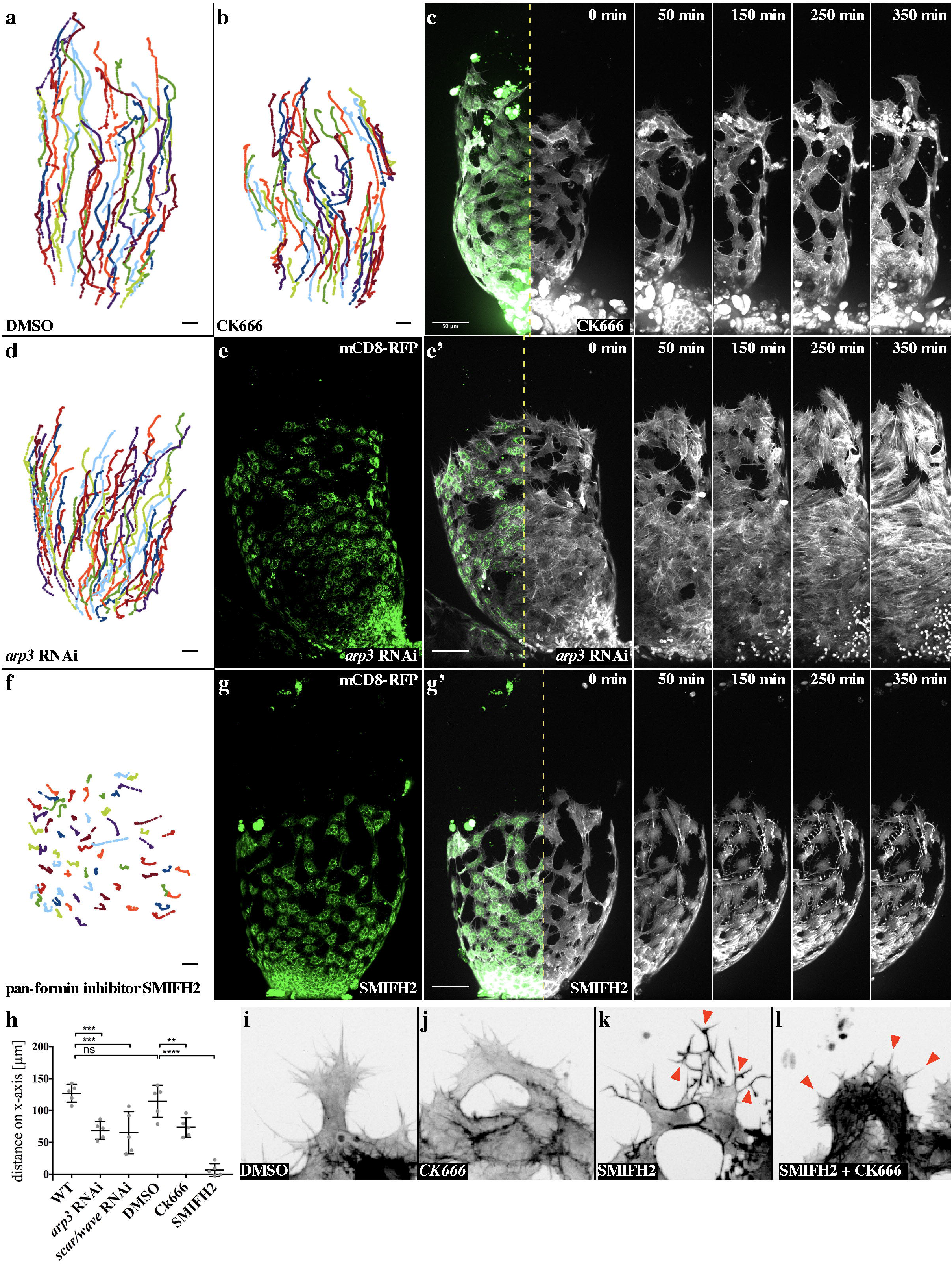
Formins are essential in myotube collective migration and filopodia dynamics, but not the Arp 2/3 complex. **a.** Migration tracks of testis myotubes 33 h APF in 420 min *ex vivo* culture, treated with DMSO as a control. Source data are provided as a Source Data file. **b, c.** CK666 (100 μM) treatment of a testis 33 h APF in 420 min *ex vivo* culture. Upon Arp2/3 complex activity inhibition, migration is reduced. Especially cells at the testis base appear to be affected. *mef2-Gal4* drives UAS-LifeAct-EGFP and UAS-mCD8-RFP expression. **b.** Migration tracks of testis myotubes upon CK666 treatment. Source data are provided as a Source Data file. **c.** Life imaging micrographs. mCD8-RFP in green and LifeAct-EGFP in white. The dashed line in c represents the area depicted in 0-420 min. Scale bar, 50 μm. **d, e.** Migration is also mildly reduced by *arp3* RNAi. *mef2-Gal4* drives UAS-LifeAct-EGFP, UAS-mCD8-RFP and the RNAi construct *UAS-arp3^KK102278^* (Vienna v108951). **d.**Migration tracks of testis myotubes upon *arp3* RNAi. **e, e’.** Life imaging micrograph. Source data are provided as a Source Data file. mCD8-RFP (green) is depicted in **e**. (Note: mononucleated myotubes marked by co-expression of the mCD8-RFP marker excluded from the nuclei). Overlay with LifeAct-EGFP (white) in **e’**. The dashed line in e’ represents the area depicted in 0-350 min. Scale bar, 50 μm. **f, g.** Upon Formin suppression through SMIFH2 (10 μM) treatment, migration is completely disrupted. *mef2-Gal4* drives UAS-LifeAct-EGFP and UAS-mcd8-RFP expression. **f.** Migration tracks of testis myotubes upon SMIFH2 treatment. Source data are provided as a Source Data file. **g, g’.** Life imaging micrographs. mCD8-RFP (green) is depicted in **g**. Overlay with LifeAct-EGFP (white) in **g’** The dashed line in **g’** represents the area depicted in 0-350 min. Scale bar, 50 μm. **h.** Quantification total migration distance along x-axis. Source data are provided as a Source Data file. **i-k.** Close ups of front-row myotubes upon different treatments. **i.** Upon DMSO treatment, cell morphology and filopodia composition were not affected. **j.** Arp2/3 suppression by CK666 treatment leads to mild defects. No branched filopodia are built, the overall morphology is unaffected. **k.** Formin suppression by SMIFH treatment leads to strong morphological defects. Cells are contracted, filopodia generate more branches. **l.** CK666 in addition to SMIFH2 co-treatment leads to a loss of branched filopodia. Cells are contracted even stronger.

By contrast, treatment with the pan-formin small-molecule inhibitor SMIFH2 ^25^ strongly affected cell morphology and completely disrupted collective myotube migration (Figure 2f, g and g’; supplementary movie M7; quantification in figure 2h, S2 g). Compared to CK666 treatment, cells treated with SMIFH2 showed a prominent reduced number of dynamic, but instead highly branched filopodia-like protrusions (Figure 2k; supplementary movie M7). Interestingly, cells co-treated with CK666 and SMIFH2 completely lacked these branched filopodial protrusions suggesting that their formation or branching depends on a still prominent Arp2/3 complex activity in SMIFH2 treated cells (Figure 2l). Supporting this notion, cells only depleted for Arp3 showed a reduction in filopodia branches resulting in a significant reduction of protrusions (Figure 3a, b; quantification in c). Consistently, an Arp3-EGFP transgene localized close to newly forming branches as we recently found in dendrite branchlet formation of *Drosophila* larval sensory neurons (arrowheads in Figure 3d, e ^26^). Interestingly, we also found a strong accumulation of the Arp3-EGFP at cell-cell contacts, an observation made in different cell systems (asterisks in Figure 3e’).

**Figure 3.**
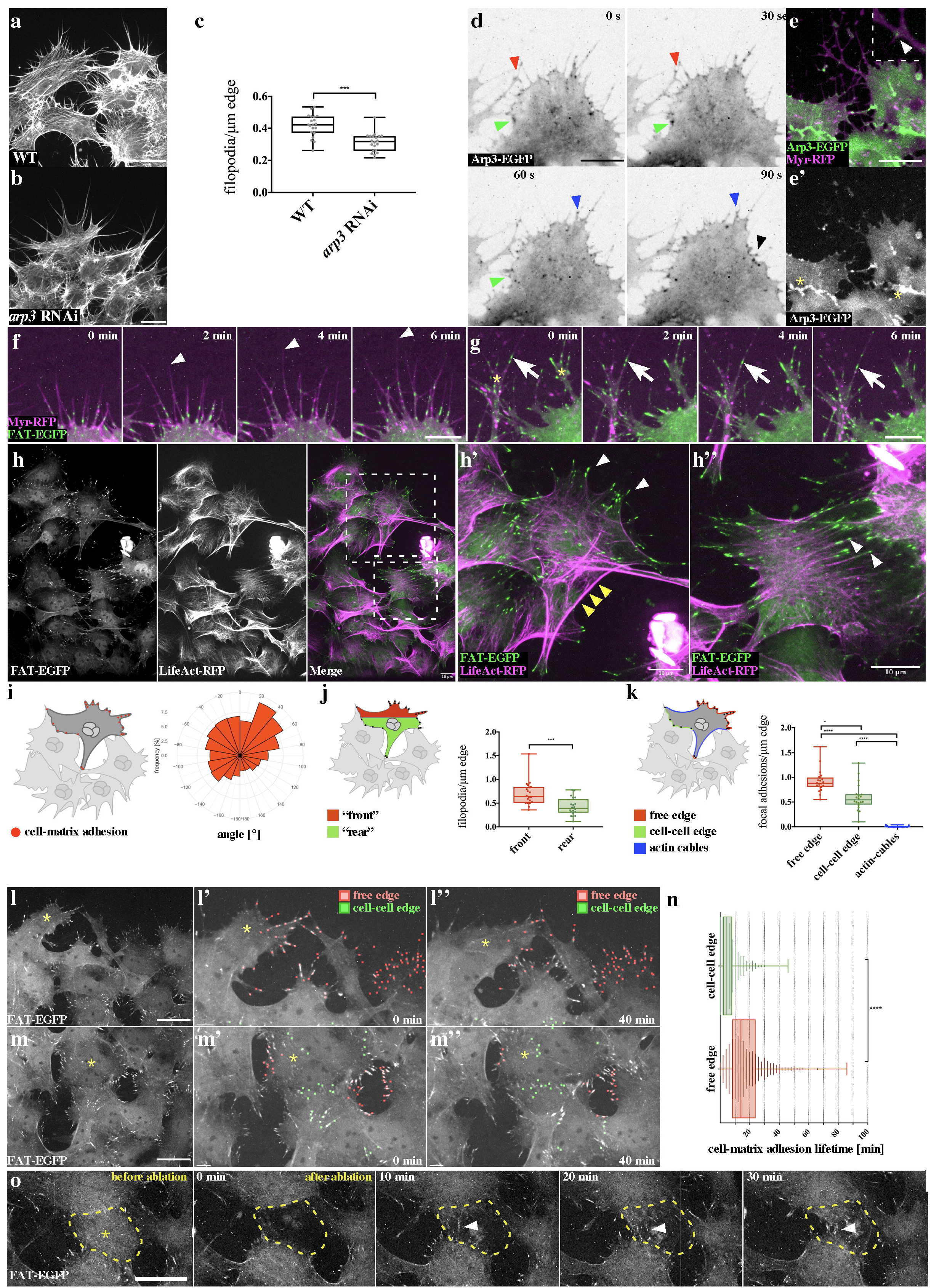
Migrating myotubes form stable cell-matrix adhesions at their free edge and adherens junctions at their cell-cell edge. **a, b.** Close ups of front-row myotubes **a.** wild type and b. *arp3* knock-down marked by Lifeact-EGFP expression. Scale bar, 10 μm. **c.** Quantification of filopodia tip number per cell edge length. Source data are provided as a Source Data file. **d.** Spinning disc microscopy still images of a front-row myotube expressing an Arp3-EGFP transgene. The arrowheads mark positions where Arp3 is enriched at filopodial branch points. Scale bar, 10 μm. **e.**Spinning disc microscopy still images of a front-row myotube co-expressing Arp3-EGFP and Myr-RFP. The arrowhead marks a position of Arp3 at a distinct filopodial branch. Scale bar, 10 μm. **f, g.** Close-up of myotubes at the front edge of the migrating sheet, 6 min in *ex vivo* culture. Cell-matrix adhesions (green) are assembled in the shafts of free edge-filopodia. Filopodia elongate, generating new adhesions in a beaded string-like manner. Scale bar: 10 μm**. g.**Some filopodia build branches. UAS-FAT-EGFP (cell-matrix adhesion marker and UAS-Myr-RFP was driven by *mef2-Gal4*. Scale bar: 10 μm. **h, h’, h’’.** FAT-EGFP and LifeAct-RFP where driven by *mef2-Gal4*. **h.** Dashed lines represent the area magnified in h’ and h’’. **h’.**Matrix adhesions are found at filopodia tips (white arrowheads) and appear enriched at the free edge of cells in contrast to their cell-cell edges. Thick actin cables are marked by yellow arrowheads. **h’’.** Cell-matrix adhesions colocalize with bundled actin fibers (white arrowheads). **h’, h’’.** Scale bar, 10 μm. **i, j, k.** Quantification revealed **i.** a significant bias in directionality of cell-matrix adhesions, **j.** an increased number of cell-matrix adhesions at the cell front (pointing to the testis apex) and **k.** an increased number of cell-matrix adhesions at the free-edge compared to cell-cell edges (excluding free edge regions with prominent actin filament bundles marked as “actin cables” in blue) as illustrated. Source data are provided as a Source Data file. **l, m.** Cell-matrix adhesions in nascent myotubes during migration. FAT-EGFP was driven by *mef2-Gal4*. Quantified matrix adhesions at the free edge are depicted in red and at the cell-cell edge in green. Scale bar: 10 μm. **l.** Front edge of the migrating sheet. **l’.** Magnification at 0 min. **J’’.** Magnification at 40 min. **m.** Following cells in the same sheet as in H. **m’.** Magnification at 0 min. **m’’**. Magnification at 40 min. **n.** The quantification revealed that cell-matrix adhesions longevity is significantly higher at the “free edge” compared to the “cell-cell-edge”. n=3 testes. Source data are provided as a Source Data file. **o.** Cell-matrix adhesions in myotubes in the middle of the migrating sheet 33 h APF in *ex vivo* culture before and after laser ablation. *mef2-Gal4* drives UAS-FAT-EGFP expression. Before ablation only few and scattered cell-matrix adhesions can be observed in “follower” myotubes. After laser ablation, cells adjacent to the ablation site start to generate cell-matrix adhesion containing protrusions along the newly arose free edge (arrowhead). Scale bar: 10 μm.

Taken together, these findings suggest that Arp2/3 activity is required in filopodia-branching, whereas the activity of formins are essential to generate filopodial protrusions. RNAi-mediated suppression of single *Drosophila* formins did not result in prominent protrusion or migration defects (see supplementary table 1), suggesting a potential redundant and synergistic functions of different formins in protrusion formation.

### Migrating myotubes preferentially form more stable cell-matrix adhesions at their free edge

It is generally believed that filopodia may promote mesenchymal migration by promoting cell-matrix adhesiveness at the leading edge to stabilize the advancing lamellipodium ^27^. Migrating myotubes lack lamellipodia, but instead filopodia appear to be critical for myotube migration as inhibition of their formation by interfering with formin function results in a complete loss of migration. Expression of a cell-matrix adhesion targeting reporter (FAT-EGFP ^28^) revealed that migrating myotubes indeed formed numerous cell-matrix anchorage sites at the base, along the shaft, and at the tip of filopodia (Figure 3f; supplementary movie M8). Multiple cell-matrix adhesions were built in a single filopodium, giving them a beaded appearance (Figure 3g; supplementary movie M8). Cell-matrix adhesions formed along filopodia shafts subsequently seemed to move rearwards, along a retrograde flow of bundled actin filaments, eventually getting disassembled in the outer rim of the cell body (Figure 3g; supplementary movie M8). Co-expression with a LifeAct-RFP reporter marked especially thicker actin bundles attached to large, more elongated cell-matrix adhesion structures that shows a more classical appearance of matrix adhesions found in lamellipodia (Figure 3h-h”).

Remarkably, the number of cell-matrix adhesions within single cells at the front edge of the cluster correlates with the presumed direction of migration towards the testis tip (Figure 3k). Cells formed an increased number of cell-matrix adhesions at the migrating front (pointing to the testis apex; figure 3i) when compared to the rear. An even more pronounced difference becomes apparent, when instead of comparing front and rear, cell-matrix adhesions are divided into those build at the free edge (excluding free edge regions comprising actin cables marked in blue) versus the cell-cell edge as illustrated in figure 3j, k. Quantitative analysis of matrix adhesion dynamics further showed that cell-matrix contacts formed at free edges showed significantly increased lifetime compared to those close to cell-cell contacts (Figure 3l-l”; supplementary movie M9; quantification in figure 3n). This asymmetric distribution of cell-matrix adhesion implies that polarization along the cell edge of myotubes does not require specialized leader cells, as observed in endothelial cells or border cell migration ^29^. It rather appears to be a response on exhibiting free edge and potentially can occur in every cell within the cluster. Consistently, an increase of free edges within the cell cluster was accompanied with the formation of new matrix adhesions as ablation experiments showed. Myotubes immediately migrated when exposed to an empty space and filled the gaps within laser-induced wounds (Figure 3o; supplementary movie M10).

### Reduced N-cadherin expression promotes single cell migration at the expense of collective directionality

Reduced cell-matrix adhesion density of myotubes in contact might be due to an enhanced disassembly of cell-matrix complexes at cell-cell contacts as previous reported for neural cells undergoing CIL ^13^. Migratory myotubes predominantly express N-cadherin as a key adhesion molecule of cell-cell contacts ^30^, which is essential in early *Drosophila* embryogenesis ^31^. N-cadherin was not only found along adjacent membranes of myotube sheets at the testis base (Figure 4a, b, b’), but were also highly enriched along the bridges of interdigitating filopodia (Figure 4c, c’, e, e’). In contrast, single myotubes without any cell neighbor that were rarely observed (Figure 4f, f’) completely lacked N-cadherin clusters at their free edge filopodia. Live imaging of migrating myotubes expressing a N-Cad-EGFP transgene confirmed a highly dynamic accumulation at cell-cell contacts and along filopodia forming initial contacts between neighboring cells (Figure 4g; see also supplementary movie M11). To further test the importance of N-cadherin-dependent cell-cell contacts in controlling the collective behavior of myotubes we used an RNAi approach to downregulate N-cadherin expression in myotubes by using the *mef*2-Gal4 driver (Figure 4h, i; supplementary movie M12). Expression of two different RNAi transgenes efficiently downregulates N-cadherin protein level as shown by immunostainings of adult testes (Supplementary figure S2 h, i, quantification in S2 j). Myotubes depleted for N-cadherin are still able to migrate, and even change more frequently their relative positions with each other within the moving cluster (Figure 4h, i; supplementary movie M12). Expression of different *N-cad* RNAi transgenes resulted in an obvious increase of free cell edges with prominent cell-matrix adhesions (Figure 4j, k; supplementary movie M13) and increased gaps between migrating myotubes (Figure 4l, m) in a dosage-dependent manner, but did not affect the cell number or cell size (Supplementary figure S2b, c, d and e). Consistently, suppression of N-cadherin led to a significantly decreased neighbor permanency suggesting that indeed a reduced N-cadherin function weakened cell-cell adhesions (Figure 4n). Quantitative analysis of the migration pattern of individual cells further revealed prominent changes of the migratory behavior. Overall, the total migration distance along the x-axis was not affected indicating that N-cadherin-depleted cells migrate as far as wild type cells (Figure 4o; supplementary movie M12). However, N-cadherin-depleted cells migrate significantly less directional but faster compared to wild type cells (Figure 4p, q). Thus, myotubes did not display a leader-follower cell dynamics, in which leader cells drag inherently passive followers cells by means of strong cell-cell cadherin contacts. By contrast, N-cadherin-mediated cell-cell contacts seem to be required for the directionally coordinated migratory behavior of myotubes.

**Figure 4.**
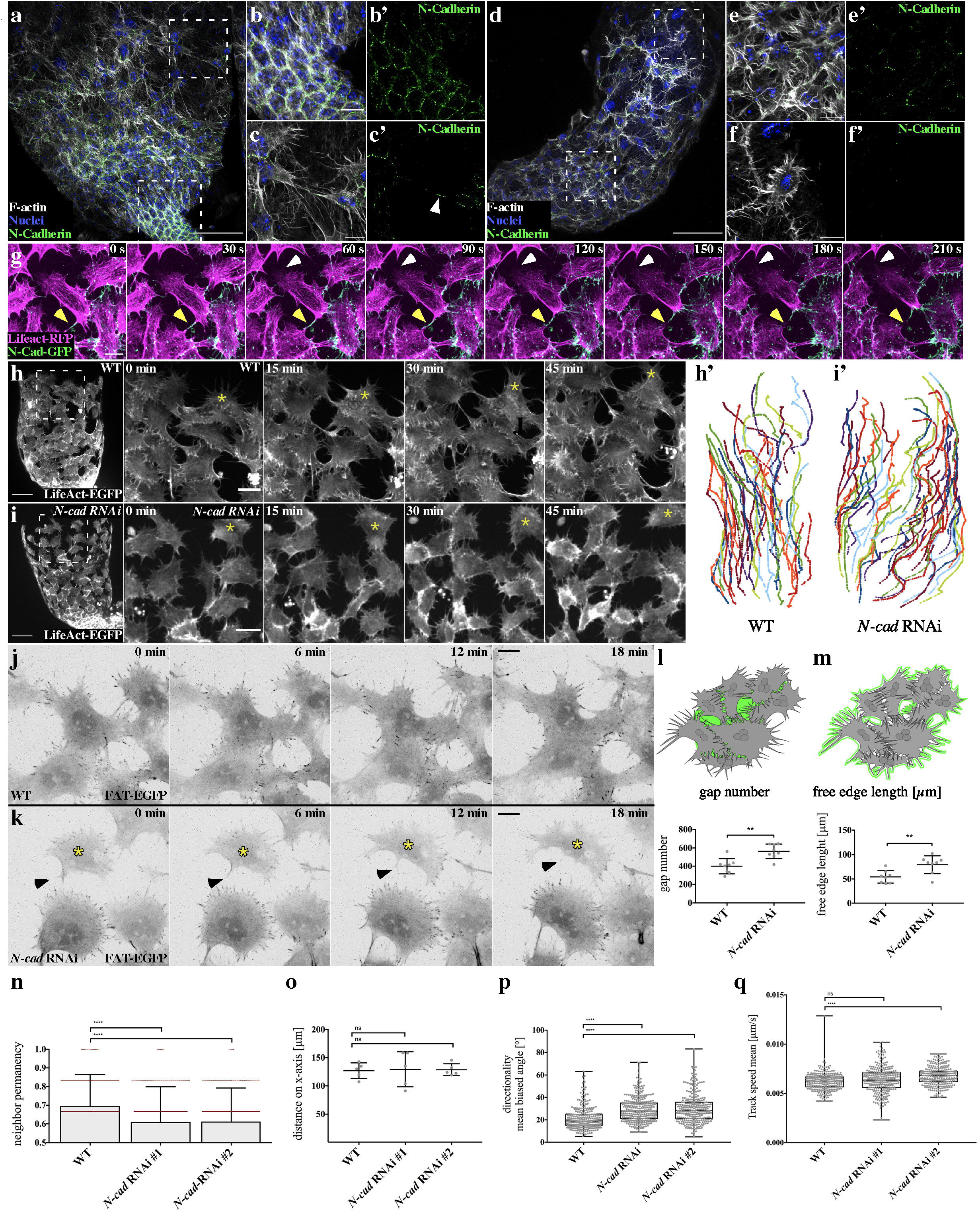
Reduced N-cadherin expression increases free edge, promoting cell-independent behavior at the expense of collective directionality. **a-f.** Confocal images of a wild type 33 h APF testis stained with an anti-N-cadherin antibody. F-Actin was stained using Phalloidin and nuclei were marked with DAPI. **a.** Overview of the testis base. The areas marked with dashed lines are magnified in B, C and E, F, respectively. Scale bar: 50 μm. **b.** On the genital disc adjacent to the testis base, nascent myotubes appear epithelial with N-cadherin localized evenly along the cell edge. **c, c’.** In contrast, at the front edge of the migrating sheet, N-cadherin localized in foci at the tip of filopodia-like structures interconnecting cells (white arrowhead). **d, e, e’.** The same is true for cells within the sheet. **f, f’.** In rare cases, completely isolated cells could be observed. No N-cadherin staining can be detected in such cells. **b, c, e, f.** scale bar: 10 μm. **g.** Spinning disc microscopy still images of myotubes expressing a Ncad-EGFP transgene. The arrowheads mark positions where Ncad-EGFP is enriched at cell-cell junctions. Scale bar, 10 μm. **h, i.** *mef2-Gal4* was used to drive expression of UAS-LifeAct-EGFP and UAS-Ncad-RNAi construct. **h**. Wild type (WT) testis 33 h APF in *ex vivo* culture. *mef2-Gal4* drives expression of UAS-LifeAct-EGFP. Overview at t = 0 min at the left side. Scale bar: 50 μm. Time steps from 0 min to 45 min in *ex vivo* culture at the right side. Scale bar: 20 μm. **h’.** Migration tracks of WT myoblasts. Source data are provided as a Source Data file. **i.** *n-cadherin* knock down in myotubes. Compare to A. Single cells are not as strongly attached to each other. Myotubes are sometimes completely isolated as in WT (yellow asterisk) (t = 0 min, 45-60 min). **i’.** Migrations tracks upon knock down of *N-cadherin*. Source data are provided as a Source Data file. **j, k.** 18 min life culture demonstrates that an increased free edge in every single cell (yellow asterisk) through N-cad RNAi results in more cell-matrix adhesion-producing filopodia. The arrowhead marks a retracting protrusion. UAS-FAT-EGFP and *UAS-Ncad* RNAi was driven by *mef2-Gal4*. Scale bar: 20 μm. **l, m**. Quantification with Fiji. Graphical representation of the values is depicted. A ROI of the same size of 8 stills of each respective genotype was compared with Fiji Particle Analysis after conversion to black-and-white pictures (see also supplementary figure S2B, C). **l.** The number of gaps between cells is significantly increased. **m.** As proxy for free edge, we used the perimeter of the white-to-black edge. Cell free edge is significantly increased in *N-cad* RNAi animals. Source data are provided as a Source Data file. **n.** Neighbour-permanency is significantly reduced when N-cadherin is knocked down (using two independent RNAi transgenes). Source data are provided as a Source Data file. **o.** To assess, how far cells were able to migrate on the testis, the difference of x values (x-axis = defined as the axis from base to apex) of testis myotubes at t = 0 min and t= 420 min was calculated. The mean of each testis was compared. Upon N-Cadherin reduction, myotubes come as far as in WT. Source data are provided as a Source Data file. **p.** As a tool for directionality, biased angle in regard to the testis axis was measured. Datasets were smoothed and Mercator-projected before (see also supplementary figure S3). The mean angle (0-180°) of every track is blotted. N-Cadherin reduction causes myotubes to migrate less directional. The same is true using a second RNAi line. Source data are provided as a Source Data file. **q.** Quantification of track speed mean in μm/sec. RNAi line #1 is subject to wider fluctuation but not significantly faster. RNAi line #2 is significantly faster than wild type (WT). Source data are provided as a Source Data file.

### Migrating myotubes need cell-cell contact to achieve directionality

To further test whether myotubes require cell-cell contacts for their directional cell migration, we performed laser ablation experiments. Isolation of single myotubes by laser ablation of the adjacent neighboring cells on the testis created a situation, in which a cell is surrounded by free edge. After the ablation, the isolated cells immediately ceased directional migratory behavior and cells formed numerous filopodial protrusions pointing in all directions (Figure 5a-c; supplementary movie M14). Once those cells got in close contact to adjacent cells, they started to migrate forward along those migratory sheets as a collective (Figure 5d, e; supplementary movie M14). Single cell tracking before and after cell-cell contact confirmed a contact-dependent migratory cell behavior of myotubes, a phenomenon that is reminiscent of CIL (Figure 5e-g). Remarkably, such a contact-stimulated migratory behavior could not be observed between two individual cells, which were still connected by cell-cell junctions but isolated from remaining cell cluster by laser ablation (Figure 5h; supplementary movie M15). Cell pairs neither migrated away from each other nor became polarized pointing protrusions into opposite directions, but instead always stuck together with constant contact distance over time (Figure 5 i, j supplementary movie M15).

**Figure 5.**
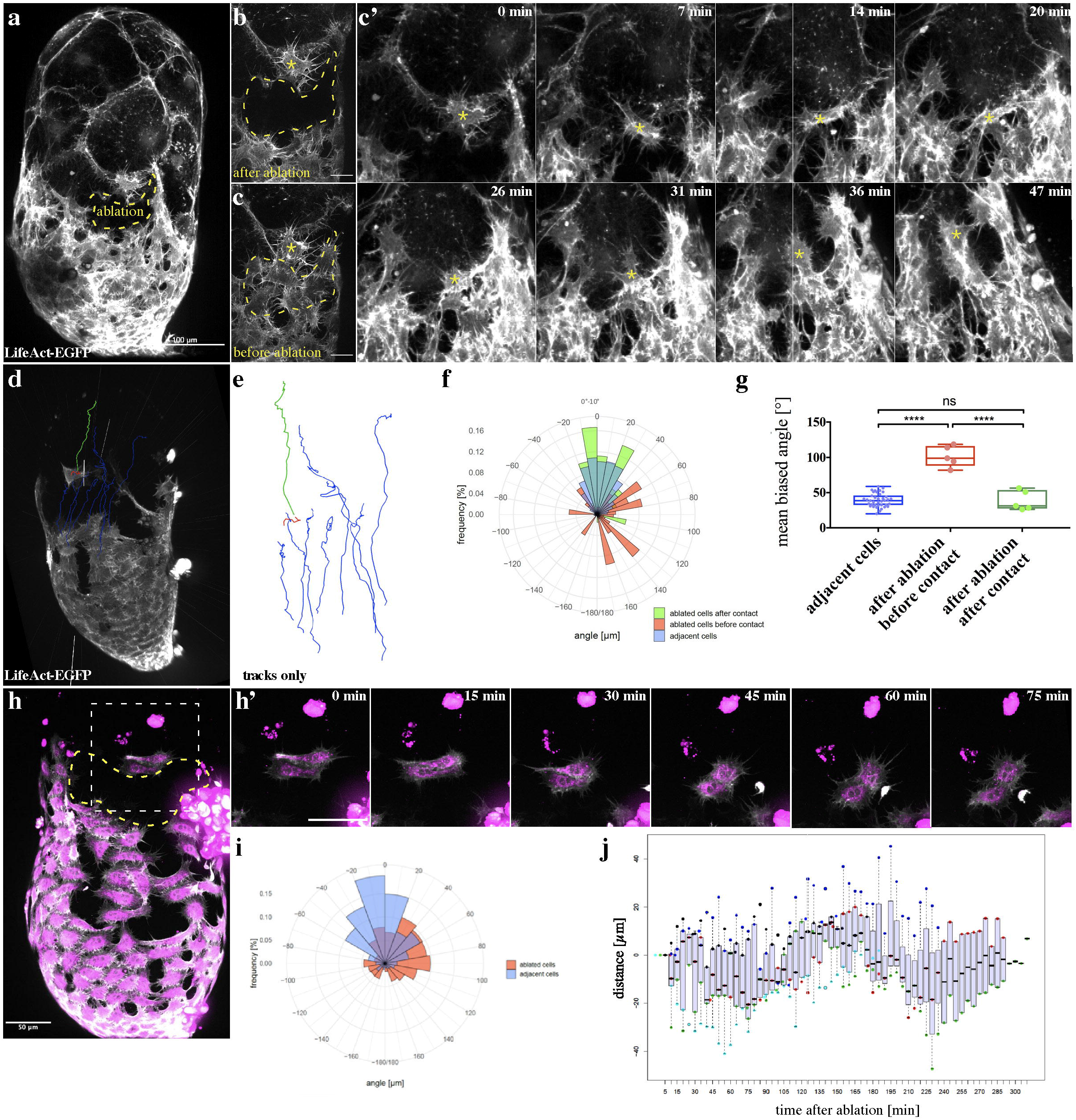
Migrating myotubes need cell-cell contact to achieve directionality. **a-g.** Isolation of a single nascent myotube by laser ablation. **a.** Overview of a testis after laser ablation (33 h APF). *htl-Gal4* drives UAS-LifeAct-EGFP expression. Scale bar, 100 μm. **b.** Close-up on the ablation site. **c.** Same site as in b, before ablation. Scale bar in c and c’: 20 μm. The dashed line represents the area affected by laser ablation. **c’.** Behavior of the isolated cell from B after ablation. The isolated cell (yellow asterisk) shows no forward motion if it has no contact to adjacent cells (upper row). After contact is established, it moves along in the migrating sheet (bottom row). **d, e.** To quantify the directionality of the isolated cell, cell motion was tracked using the Imaris software. The isolated cell before contacting to the migrating sheet is depicted in red, after contacting it is depicted in green. As a control, adjacent cells were tracked. They are showed in blue. Source data are provided as a Source Data file. **f, g.** As a measurement tool, we used the biased angle to x-axis. The mean angle (0-180°) of every track is blotted. When isolated, cells lose their directionality, but regain it after establishing contact to adjacent cells. The color code is the same as in **e, f**. n= 5 testes. Source data are provided as a Source Data file. **h-j.** Isolation of two adjacent myotubes by laser ablation. **h.** Overview of a testis after laser ablation (33 h APF). *htl-Gal4* drives LifeAct-EGFP (grey) and Myr-RFP (magenta) expression. The dashed line represents the area affected by laser ablation. Scale bar, 100 μm. **h’.** Behavior of the two isolated myotubes from h. after ablation. Scale bar in c and c’: 20 μm. **i.** Rose plot shows the distribution of the biased angle to x-axis. Source data are provided as a Source Data file. **j**. Measurement of the distance between two myotubes over time. Source data are provided as a Source Data file.

### Rac2 and Cdc42 functions play important roles in myotube migration shaping testis morphology

Cell adhesions are not only required to mechanically couple cells within the cluster, but also to link adhesion complexes to the actin cytoskeleton controlling the protrusion dynamics and directionality^32^. Rho GTPases are critical molecular players that regulate adhesions and motility during single and collective cell migration ^33,34^. To identify such key players contributing to myotube migration we used an RNAi approach to screen numerous candidate genes (see supplementary table 1). Defects in testis myotube migration during pupal metamorphosis can be identified by a prominent disturbed morphology of adult testis (supplementary figure S1a-l; ^30^. The adult testis is a pair of thin tubules of 2.5 coils and ~2 mm in length surrounded by a sheath of multinuclear smooth-like muscles ^19,30^. Defective N-cadherin-mediated cell-cell adhesion resulted characteristic holes in the muscle sheet ^19^, where myotubes were not properly attached to one another (supplementary figure S1a, c, e). In contrast, defects in myotube migration resulted in an abnormal testis morphology with reduced coils and bulky tips (Supplementary figure S1a, f-k). Depending on the phenotypic strength the muscle sheath only partially or completely failed to cover the entire testis resulting into strong elongation/coiling defects (supplementary figure S1a). Strong abnormalities were observed following RNAi-mediated suppression of Cdc42 and Rac2 functions, one of the two very similar *rac* genes in *Drosophila* ^35^. In both cases, the adult testes were smaller than in the wild type with reduced coils and bulky tips (Supplementary figure S1f, g). The muscle sheath either did not cover the entire testes with numerous large holes. In comparison, suppression of Arp2/3 complex subunits and single subunits of the WAVE regulatory complex (WRC ^36^) such as WAVE and the Rac-effector Sra-1, resulted into more moderate morphological defects compared to *rac2* or *cdc42* depletion. Adult testes deficient for Arp3, WAVE and Sra-1 still had about 1.5 to 2 coils, however many myotubes also did not reach the testis apex resulting into bulky tips (Supplementary figure S1h-j).

### Rac2 and Cdc42 are required for myotube migration by differentially regulating cell-matrix adhesions

Compared to suppression of the Arp2/3-WRC pathway, knockdown of Rac2 functions led to stronger defects in membrane protrusions and cell migration suggesting that Rac2 might have additional roles in myotube migration (Figure 6a, d; compare quantification in supplementary figure S2g). *rac2*-depleted cells showed a severely changed cell morphology with thinner and highly dynamic filopodial protrusions. These filopodia were unable to adhere stably (Figure 6g, g’; supplementary movie M16, M17). Supporting this notion, live-cell imaging of *rac2* knockdown cells using the FAT-EGFP reporter revealed a prominent loss of cell-matrix adhesion contacts (Figure 6l, supplementary movie M18). Since *mef2*-Gal4 driven FAT-EGFP is still normally enriched in integrin-dependent adhesion structures such as muscle attachment sites of the larval body wall musculature, a general impact of Rac2 function on matrix adhesion can be excluded (Supplementary figure S1m, n).

**Figure 6.**
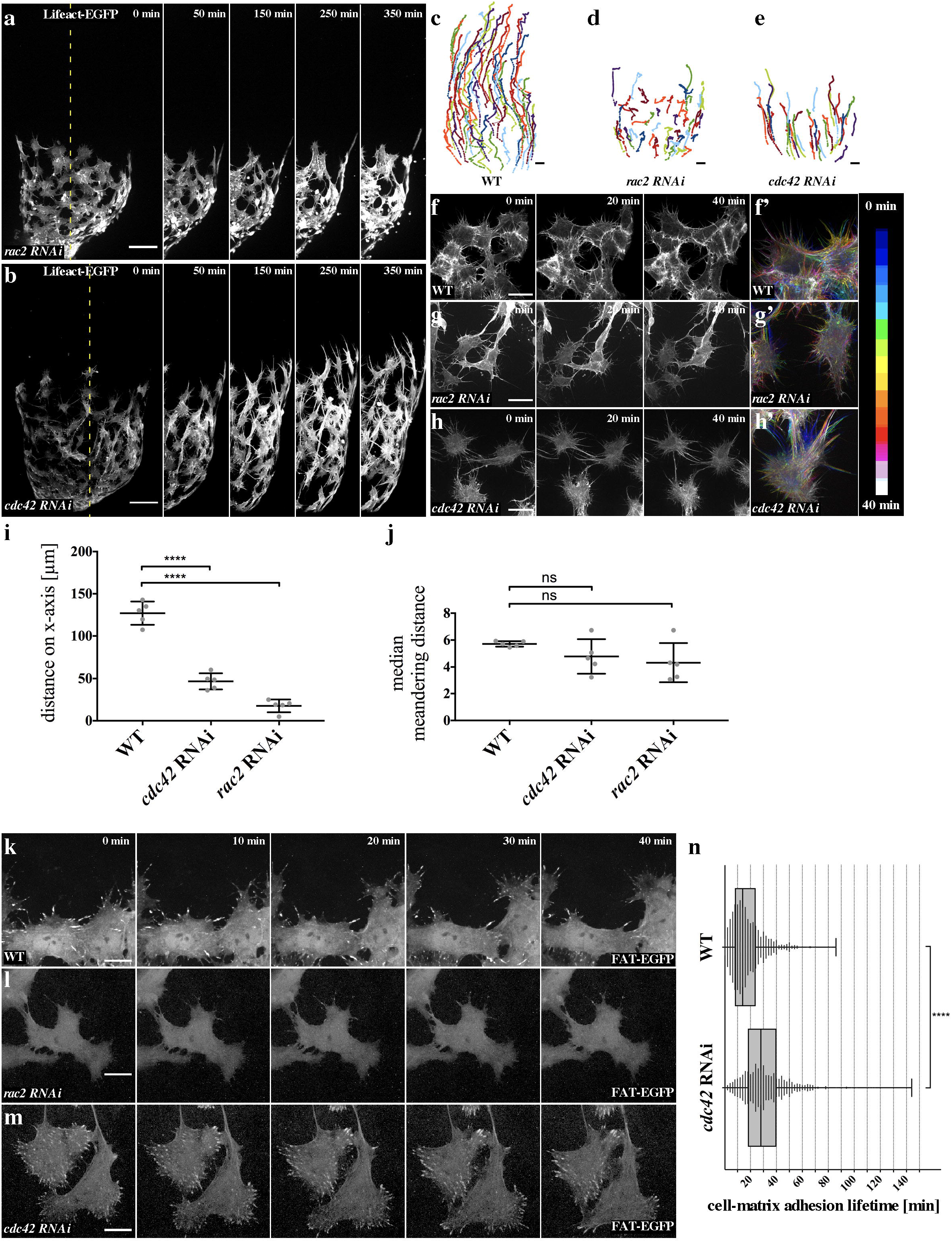
Rac2 and Cdc42 regulate filopodia matrix adhesion to enable myotube collective migration. **a, c, i, j.** *rac2* knockdown was induced by expression of the *UAS-rac2^NIG.8556R^* RNAi transgene together with UAS-LifeAct-EGFP, using *mef2-Gal4.* **a.** *rac2* knockdown in myotubes on testis 33 h APF in *ex vivo* culture. Myotube migration almost completely ceases. Tracks are depicted in D. The dashed line in “0 min” represents the area depicted in 50–420 min. Scale bar: 50 μm. Source data are provided as a Source Data file. **b, d, i, j.** *cdc42* knockdown was induced by expression of the *UAS-cdc42^TRiP.JF02855^* (#1) or the *UAS-cdc42*^*KK108698*^ (#2) RNAi transgenes, together with UAS-LifeAct-EGFP, using *mef2-Gal4.* Source data are provided as a Source Data file. **b.** *cdc42* knock down in myotubes on testis 33 h APF in *ex vivo* culture. Myotube migration is disrupted. Cells change their shape, generating massive filopodia-like structures, in comparison to WT. Tracks are depicted in **e**. The dashed line in “0 min” represents the area depicted in 50–420 min. Scale bar: 50 μm. Source data are provided as a Source Data file. **f-h.** Close-up of myotubes 33 h APF in *ex vivo* culture with corresponding color-coded projection in **f’-h’**. Scale bar: 10 μm**. f, f’.** Wild type (WT) myotubes. **g, g’.** *rac2* RNAi causes a fast assembly and disassembly of filopodia. **h, h’’.** *cdc42* RNAi leads to very stable filopodia in comparison to wt. Filopodia are prolonged, even between nascent myotubes, rendering close cell-cell contact harder to achieve, thus the entire sheet appears less dense as in WT. **i.** Quantification of migration distance on x-axis (compare to Fig 3J). Source data are provided as a Source Data file. **j.** Quantification of median meandering distance. Source data are provided as a Source Data file. **k.** Cell-matrix adhesions in myotubes during migration. UAS-FAT-EGFP was driven by *mef2-Gal4*. Scale bar: 10 μm**. l.** Cell-matrix adhesions are completely lost upon *rac2* suppression by RNAi. **m.** Cell-matrix adhesions remain much longer upon *cdc42* reduction, even reaching the trailing end of a migrating cell. **n.** Quantification of cell-matrix adhesion lifetime. As shown in H, a cell-cell edge cannot be clearly defined upon knock down of *cdc42*, thus just the free edge was compared. *cdc42* reduction increased lifetime of cell-matrix adhesions significantly, compared to WT, n = 3 testes for WT and *cdc42* RNAi. Source data are provided as a Source Data file.

Suppression of Cdc42 function also severely impaired migration speed resulting in a strongly reduced migration distance on the x-axis (Figure 6b, e; supplementary movie M15; compare quantification in supplementary figure S2g). However, compared to *rac2*-depleted cells, *cdc42*-deficient myotubes showed an increase of thin and prolonged filopodia (Figure 6g, g’; supplementary movie M17, M19). Overall, the *cdc42*-depleted myotubes showed an elongated cell shape with numerous gaps between adjacent cells. Live-cell imaging of *cdc42* knockdown cells using the FAT-EGFP reporter revealed a significantly increased lifetime of cell-matrix adhesions (Figure 6m; quantification in figure 6n; supplementary movie M18). Compared to wild type cells, the cell-matrix adhesions remained much longer, even when they reached the trailing end of a migrating cell (Supplementary movie M18). In summary, Rac2 and Cdc42 are both required for myotube migration, but appear to differentially regulate cell-matrix adhesions.

### Activated Rho1 is not enriched at cell-cell contacts

The activity of Rho1^37,38^, the *Drosophila* homologue of RhoA, appears to be as essential for myotube migration as Cdc42 and Rac2. RNAi-mediated suppression of Rho1 but not RhoL activity in myotubes indeed resulted in strong morphological defects of the testes, and even under low RNAi transgene expression (using *lbe*-Gal4 driver) *rho1* depleted myotubes showed strong migration defects (see supplementary figure S1k, table 1). Suppression of the same RNAi transgenes using the mef4-Gal4 driver resulted into an early pupal lethality (data not shown, supplementary table 1). Different from neural crest cells undergoing CIL, activated Rho1 was not enriched at cell-cell contacts between myotubes (Figure 7a). Live imaging of migrating myotubes coexpressing a Rho1-GTP biosensor or Anillin Rho-binding domain fused to GFP (Anil.RBD-GFP ^39^ and a LifeAct-RFP transgene uncovered highly dynamic, local pulses of Rho1 activity along retracting filopodial protrusions at free edges (Figure 7a, b; supplementary Movie M20). Rho1 activation appeared to be synchronous with backward movement of retracting filopodial protrusions (Figure 7b; supplementary Movie M20). Once a protrusion has been completely retracted, activated Rho1 disappeared. Remarkably, retracting protrusions were often followed by new forward-directed protrusions at the same region without any Rho1 signal (Figure 7b, Supplementary Movie M20). Thus, migrating myotubes are not simply polarized along a front-rear axis.

### Myotube migration requires Rok-dependent actomyosin contractility

Rho1 is known to control myosin II-dependent contraction through the protein kinase Rok shaping cells into tissue in a large variety of morphogenetic events during development ^40,41^. To test whether Rok-dependent actomyosin-mediated contractility is required for myotube collective migration, we first inhibited contractility by treating ex vivo cultured pupal testes with the specific Rok inhibitor Y-27632^42^ and with blebbistatin ^43^ or rather its photostable derivate para-nitro-blebbistatin ^44^, which targets the action of the myosin II (Figure 7c, Supplementary Movie M21). Compared to control cells treated with DMSO, we found similar striking changes in cell morphology in a time-dependent manner that eventually disturb myotube cell migration (Figure 7c, e, f; supplementary Movie M21). Following treatment with Y-27632 or blebbistatin, the myotube cell cluster was still able migrate, but became dramatically elongated with long interconnecting cell processes as expected for a tissue under stretch. As consequence, the cell cluster showed large gaps between individual cells, which dramatically increased in the total size over time (see quantification in figure 7h). Consistently, RNAi-mediated depletion of both the regulatory light chain of the myosin II (*spaghetti squash*, *sqh*) and the myosin II heavy chain (*zipper, zip*) phenocopies the pharmocological inhibition of Rok (Figure 7d, g, supplementary Movie M21; quantification in figure 7h). Migratory defects and the inability to tighten up the cell cluster finally led to small adult testes with reduced coils and bulky tips with numerous large holes in the muscle sheet similar to those depleted of Rho1 (compare figure 7k, l with supplementary figure S1b, k). In conclusion, these data show an important role of Rok-driven actomyosin contractility in collective myotube migration. Together, our data do not support that actomyosin-dependent contractility is required for myotube forward movement, but rather contribute to the integrity of the migrating cell cluster.

**Figure 7.**
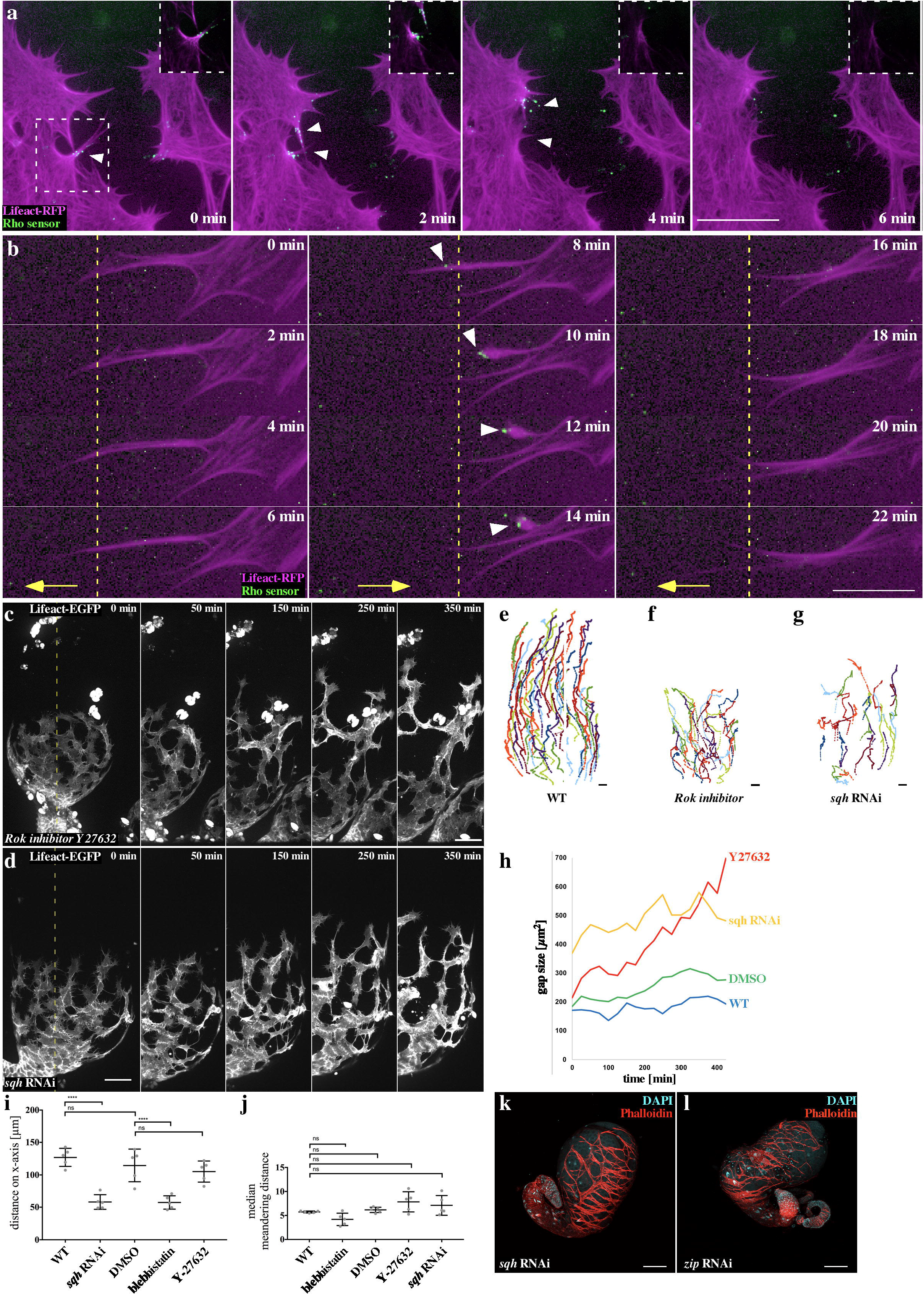
Rho/Rok-driven actomyosin contractility is essential for myotube migration. **a, b.** Close-ups of myotubes at the front edge of the migrating sheet 30 min in *ex vivo* culture. *mef2-Gal4* drives UAS-LifeAct-RFP and the Rho1 sensor Anillin-RBD-EGFP. UAS-LifeAct-RFP is enriched along the membrane in actin cables. To depict all actin structures, gamma was set on 0.09. In the boxes in the upper right corner, details with gamma=1 are depicted. **a.** Rho1-Sensor activity is found in free edge filopodia (white arrowheads, as LifeAct-RFP rapidly bleached out in filopodia tips, EGFP signal appears partially outside the cell). When analyzed with gamma=1, it becomes clear that Rho1-sensor is only present at parts of the edge containing actin cables. After the Rho signal appears, the corresponding part of the cell retracts, and the Rho1 signal immediately disappears. During retraction, the LifeAct-RFP signal at the retractive site goes back to normal intensity. **b.** Rho1 sensor activity does not seem to mark rear polarity. Filopodia can protrude (left column, yellow line, then activate RhoA and retract (middle column, yellow line). Subsequently, neighboring filopodia can elongate again (middle and left column, yellow line). **c.** myotubes expressing LifeAct-EGFP on testis 33 h APF in *ex vivo* culture treated with the Rok inhibitor Y-27632. **d.** *sqh* knockdown was induced by expression of the *UAS-sqh* RNAi transgene together with UAS-LifeAct-EGFP using *mef2-Gal4.* Myotube cell cluster were still able migrate with reduced speed and become dramatically elongated with long interconnecting cell processes. The dashed line in “0 min” represents the area depicted in 50–420 min. Scale bar: 50 μm. Tracks are depicted in **e.** wild type (WT), **f.** Y-27632 treatment and **g.** *sqh* knockdown. Source data are provided as a Source Data file. **h**. Measurement of the gap size within cell cluster over time. Source data are provided as a Source Data file. **i.** Quantification of migration distance on x-axis. Source data are provided as a Source Data file. **j.** Quantification of the median meandering distance. Source data are provided as a Source Data file. **k, l.** Confocal images of adult testes expressing **k.** a *sqh* RNAi transgene **l.** a *zip* RNAi transgene under the *mef2-Gal4* driver, the muscle sheet is stained with phalloidin (red) and nuclei are stained with DAPI (cyan). Scale bar: 100 μm.

## Discussion

### Myotube migration – a new model system for collective cell migration

In this study, we established a new model system for studying collective cell migration in organ culture that allows high-resolution long-term live-imaging microscopy combined with genetic, pharmacological, and mechanical perturbation analysis. Our data implies that a contact-dependent migration mechanism acts as a driving force to polarize *Drosophila* myotubes and to promote their directional movement along the testes. A contact-stimulated migration has been already observed in cultured cells many years ago, but the molecular mechanisms underlying this phenomenon has been never analyzed in more detail ^45^. Thomas and Yamada observed that both primary neural crest cells and two neural-crest-derived cell lines barely moved when isolated in suspension, but could be stimulated up to 200-fold to migrate following contact with migrating cells ^45^. This process might help to ensure the cohesion and coordination of collectively migrating myotubes to form dense muscular sheets in the walls of developing hollow organs. Those muscle fibers that race ahead will immediately cease migration when they lose contact with their neighbors. That is exactly what we observed in our experiments. After ablation, an isolated myotube awaits restimulation by the other cells of the migrating cluster. Consistently, reduced N-cadherin function promotes single cell migration toward the free space at the expense of collective directionality. The contact-dependent behavior of myotubes also resembles contact inhibition of locomotion (CIL), a well-characterized phenomenon ^16^. CIL regulates the *in vivo* collective cell migration of mesenchymal cells such as neural crest cells by inhibiting protrusions forming within the cluster at cell-cell edges and by driving actin polymerization at their free edge ^46^. Different from neural crest cells, myotubes did not migrate as loose cohorts, but maintain cohesiveness (see model in figure 8). In the context of more-adhesive cells, a CIL-related mechanism, termed “frustrated” CIL has been proposed by which cell-cell junctions can determine the molecular polarity of a collectively migrating epithelial sheet ^47,48^. The authors provided evidence that cell-cell junctions determine the molecular polarity through a network of downstream effectors that independently control Rac activity at the cell free end and Rho-dependent myosin II light chain (MLC) activation at cell-cell junctions ^47,48^. At the first glance myotubes do not show an obvious polarized cell morphology with prominent polarized protrusions. Instead, myotubes form numerous competing protrusions in all directions. However, protrusions pointing to the free space preferentially form more stable cell-matrix adhesions as anchorage sites for forward protrusions, whereas the lifetime of cell-matrix adhesions at cell-cell contacts is decreased. Thus, a contact-dependent asymmetry in matrix adhesion dynamics seems to be important for the directionality of migrating myotubes, a molecular polarity that has been also found in neural crest cells undergoing CIL ^13,49^. Only when one of the adhesions of competing protrusions disassembles, pulling of the cell body towards the competing protrusions might contribute to symmetry breaking and directionality of collective migration (see model in figure 8).

**Figure 8.**
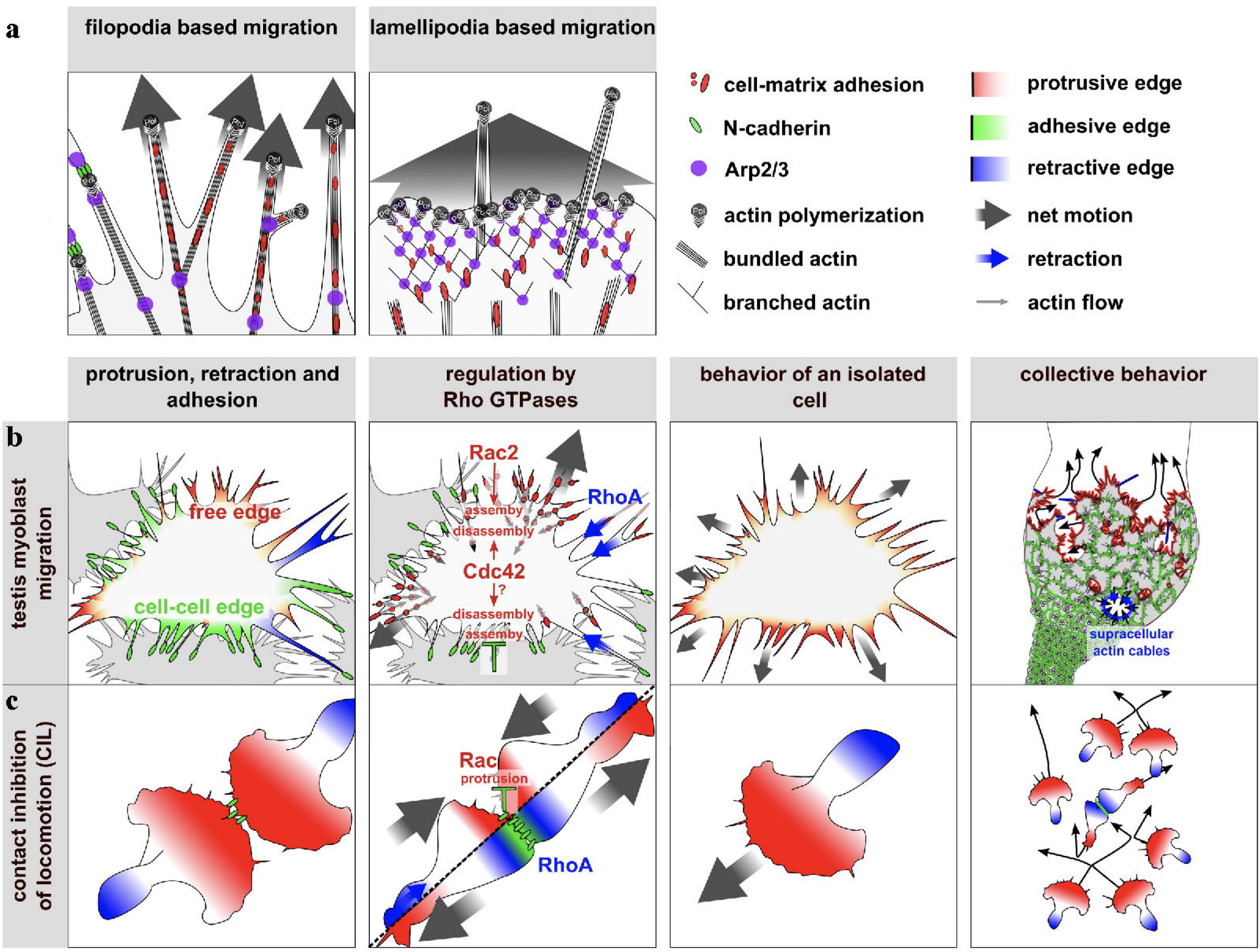
Proposed model. **a.** Comparison between filopodia-based and lamellipodia-based cell migration. Lamellipodia-based migration requires the Arp2/3 complex generating branched actin filament networks that serve as the major engine to push the leading edge forward, whereas filopodia support mesenchymal migration by promoting cell-matrix adhesiveness at the leading edge stabilizing the advancing lamellipodium or by sensing the environment. In filopodia-based migration, it seems that filopodia replace the lamellipodium as the motor of motility. We assume that polymerization of bundled actin filaments through formins pushes parts of the membrane. Arp2/3 complex contributes to filopodia branching and thereby provides new barbed ends generating new filopodia. **b.** Key features and cell behavior in testis myotube migration compared to **c.** migrating mesenchymal neural crest cells undergoing CIL. Different from neural crest cells, myotubes did not migrate as loose cohorts, but maintain cohesiveness. Unlike neural crest cells, migrating myotubes are not simply polarized along a front-rear axis and do not form a contact-dependent intracellular Rho gradient that initiates cell polarization driving directed cell migration. In myotube migration, a contact-dependent asymmetry of cell-matrix adhesion rather acts as a major switch to drive locomotion towards the free space. Individually or loosely connected migrating cells, like neural crests cells are able to migrate persistently due to classical front-rear polarity. By contrast, testis myotubes rely on constant cohesion to break symmetry. Supracellular contractile actin cables contribute to the integrity of the migrating cell cluster and thereby to cohesion.

### Rho GTPases differentially regulate myotube migration

We further provide evidence for a differential requirement of the Rho GTPases, Rac2 and Cdc42 in regulating cell-matrix adhesion. *cdc42* knockdown cells formed less cohesive clusters and showed a significant increase of cell-matrix adhesion lifetime probably due to a decrease cell-matrix adhesion turnover. In contrast, Rac2 depletion resulted in a prominent loss of cell-matrix adhesions, a phenotype that has already been described in Rac1^-/-^ mouse embryonic fibroblasts ^50^. Thus, we propose a model in which cell-matrix adhesions are downregulated at N-cadherin-dependent cell-cell contacts, a process that requires Cdc42 functions. To finally test whether a contact dependent reduction of cell-matrix adhesion in filopodia is sufficient to explain the observed collective cell behavior, we developed a simplified simulation model with a few rules governing cell behavior such as protrusive filopodia, matrix adhesion, cell-cell adhesion, and membrane resistance (Supplementary movie M22). Unlike comparable computer models ^51,52^, single cells do not possess directional information. A cell’s position is defined by the geometric center of all its filopodia, whose emergence/disappearance/elongation causes translation of the centroid, perceived as motion. Upon cell-cell contact, filopodia lose their cell-matrix adhesion and thereby their grip on the ECM, but keep connections through cell-cell adhesions. These adhesions are recognized by both contributing cells to calculate their respective centroids (see supplementary material). Using these simple rules, we could indeed model myotube collective migration, provided that cells are positioned in a confined area mimicking the unfolded testis surface (Supplementary movie M22 2A, 2B). If filopodia disappear directly after contact, cells exhibit a different cell behavior that is very reminiscent of CIL (Supplementary movie M22 2C). This simplified model further confirms our observation that local regulation of cell-matrix adhesion suffices to drive collective motility.

### Actomyosin function ensures the integrity of cohesive myotube cluster during migration

Myotube migration also requires Rho1 the *Drosophila* homologue of RhoA. Different from cells undergoing CIL, in migrating myotubes activated Rho1 was not enriched at cell-cell contacts between myotubes, but rather localized as local pulses along retracting filopodial protrusions at free edges. The effects of tensile forces have to be addressed separately in the future, by establishing one of the many existing force measurement techniques such as transition force microscopy (TFM) or using *in vivo* FRET-based tensions sensors in this system. We show that loss of Rok activity, *sqh* and *zip* phenocopies *rho*1 knockdown suggesting that a canonical pathway controls myotube migration in which Rho1 acts through Rok kinase to activate myosin II contractility. This finding supports the notion that in testis myotubes, unlike many other cell types, locally restricted Rho-GTPase regulation outweighs global Rac/Rho regulation along the cell-rear axis to achieve directionality. Previous studies demonstrated that myosin II-dependent contraction is essential for coordinating the CIL response in colliding cells. In myotube migration, Rok-dependent actomyosin contraction seems to be not required to drive the myotube cluster forward, but rather contractile actin cables contribute to the integrity of the migrating cell cluster. Thus, myotube cluster behave more like a collectively migrating monolayered epithelial sheet during gap closure^53^. While myotubes migrate into any given free space, they leave larger gaps within the cell sheet surrounded by prominent circumferential actin cables. Constriction of these supracellular actin cables necessarily might lead to gap closure observed in wildtype, but not in cells defective for RhoRok-driven actomyosin contractility.

### Filopodia based-myotube migration depends on the differential function of formins and the Arp2/3 complex

Efficient mesenchymal cell migration on two-dimensional surfaces is thought to^54^ require the Arp2/3 complex generating lamellipodial branched actin filament networks that serve a major engine to push the leading edge forward.

Interestingly, epithelial and mesenchymal cells form more filopodia when the Arp2/3 complex is absent ^55–57^. Under these conditions, mesenchymal cells lack lamellipodia and adopt a different mode of migration only using matrix-anchored filopodial protrusions. Our data further provide evidence for a filopodia-based cell migration in a physiological context during morphogenesis. This migration mode largely depends on formin as central known actin nucleators generating filopodia ^33,58^. Our data also suggest that the Arp2/3 and its activator, the WRC contribute to a more efficient myotube migration by promoting filopodia branching, and thereby increasing the number of cell-matrix adhesions, thus increased anchorage sites. Overall, filopodia-based migration enables the cell to regulate discrete subunits of membrane protrusions as an answer to the environment. The sum of filopodial protrusions adds up to a net cell locomotion that occurs similarly during lamellipodial migration please compare figure 8). Filopodial matrix adhesion complexes not only provide anchorage sites, but also allow cells to directly restructure their microenvironment by membrane-bound matrix proteases. There is indeed increasing clinical evidence suggesting filopodia play a central role in tumor invasion ^27,59^. Similar to invading cancer cells myotubes rather migrate through a 3D microenvironment composed of extracellular matrix restricted by pigment cells from the outside of the testis. Thus, it will be interesting to determine to what extent extracellular matrix restructuring by metalloproteinases is required for myotube migration.

Taken together, our data suggest that contact-stimulated filopodia-based collective migration of myotubes depends on a CIL-related phenomenon combining features and molecular mechanisms described in mesenchymal and epithelial sheet migration as well. We propose a model in which contact-dependent asymmetry of cell-matrix adhesion acts as a major switch to drive directional motion towards the free space, whereas contractile actin cables contribute to the integrity of the migrating cell cluster.

## Experimental procedures

### Drosophila Genetics

Fly husbandry and crossing were carried out according to the standard methods ^60^. Crossings and all UAS-Gal4-based Experiments including RNAi were performed at 25 °C. The following fly lines were used: *mef2*-Gal4 ^61^, *beatVC*-Gal4 (BL-40654), *htl*-Gal4 (BL-40669), *lbe*-Gal4 (BL-47974), UAS-LifeAct-EGFP (BL-35544), UAS-LifeAct-RFP (BL-58715), UAS-GFP nls (BL-4775), UAS-mcd8-RFP (BL-32219), UAS-myr-mRFP (BL-7119), cell-matrix adhesion sensor UAS-fat-GFP ^28^; RhoA-activity sensor Ubi-Anillin.RBD-GFP ^39^. All UAS-RNAi lines we used are summarized in table 1 (Supplementary data). CyO/Sco; TM2/TM6B was used as a tool for multi-step crossings, control crossings were conducted using *w*^*1118*^.

### Immunohistochemistry and fluorescence staining

Adult and pupal testis fixation and antibody staining was performed as described elsewhere ^19^. The following antibody was used: anti-Cadherin-N (1:500, DSHB DN-Ex #8). The following secondary antibodies were used: Alexa Fluor 488 (*Molecular Probes*). Alexa Fluor Phalloidin 568 (*Molecular Probes*) staining on pupal testes was carried out during the secondary antibody incubation for 2 h (1:1000 in PBS). Adult testes were stained overnight (1:1000 in PBS). DAPI (*Molecular Probes*) was performed for 10 min.

### Microscopy/4D live cell imaging of testicular nascent myotubes

Fixed pupal testes were embedded in Fluoromount-G (SouthernBiotech) and imaged on object slides. Adult testes were imaged in live-culture dishes in PBS, to maintain their natural shape. Light micrographs were taken with a Leica M165 FC stereo microscope equipped with a Leica DFC7000 T CCD camera. All fluorescent microscopic stills were taken with a Leica TCS SP8 with a HC PL APO CS2 20x/0.75 dry objective. 4D live cell imaging was performed on developing testes of 33 h APF pupae. Prepupae were collected and timed as described elsewhere ^30^. Life imaging of pupal testes was performed like on egg chambers, as described before ^62^. Images were taken on a Zeiss Observer.Z1 with a Yokogawa CSU-X1 spinning disc scanning unit and an Axiocam MRm CCD camera (6.45 μm x 6.45 μm). Long-term imaging was performed using a LD LCI Plan-Apochromat 25x/0.8 Imm Korr DIC oil-immersion objective over 7 h, with a z-stack every 5 min. Close-ups were taken with a C Plan-Apochromat 63x/1.4 oil-immersion objective over 2 h, with a z-stack every 2 min. Laser ablation of single cells on the testis was performed with a Rapp TB 355 laser.

### Chemical inhibitors

Live imaging experiments with chemical inhibitors were performed exactly as described above. All inhibitors were pre-solved in DMSO and stored at −20°C. The following inhibitors were used: CK666 (100 μM, *Sigma-Aldrich*), Formin inhibitor SMIFH2 (10 μM, *Abcam*), para-nitro-blebbistatin (10 μM, Cayman Chemical), Rok inhibitor Y-27632 (10 μM, Cayman Chemical).

### Data processing and quantification with Fiji

Filopodia angles were obtained by manually tracking filopodia tips using the *Multiple Points* tool. The center of mass was calculated in Fiji. Testes for single cell analysis (marked with beatVC-Gal4 ≫ lifeact-EGFP) were always oriented with the testis tip, the presumptive destination, pointing left (See Figure 1m’). The angle of the vector between filopodia tip and center of mass was calculated in R using the package *matlib*. Rose plots where generated using the package *ggplot2*. Membrane length for filopodia density (number per μm membrane) was quantified with the *Free Hand Line* tool and R. For single cell analysis of oriented images, points left of a virtual horizontal line crossing the center of mass (front) where compared to points on the right-hand side (rear, see Figure 1k). For processing and quantification of adhesion defects on still images a ROI with a defined size (120 × 220 px), in the middle of the migrating sheet was chosen. To obtain a black-and-white image for further analysis, a threshold was set (min: 299, max: 300). The cell number inside the ROI was counted. All further values were assessed using the *Analyze>Analyze Particles*-Tool. Area per cell was derived from the total area/cell number. Free cell edge was derived from the sum of all perimeters, as they constitute the length of black-to-white border, which is tantamount to the free cell edge. The gap number was derived from the number of coherent particles, when black-and-white picture are inverted (see also supplementary figure S2 B, C). The size of gaps during life imaging was measured with *Analyze>Analyze Particles*-Tool, too. Only gaps larger than 20 μm^2^ where analyzed. Corrected total cell fluorescence (CTCF) was measured on sum-projections based on the method established elsewhere ^63^.

### Data processing and quantification of 4D life image stacks

Manual tracking of migrating myotubes was performed using the *spots*-module in the Imaris 9.3 software. For drift correction, the *reference frame* module was used. The x-axis was positioned as axis from the genital disc to the testis hub. Excel was used for all processing and quantification. ***Distance on* X** is defined as the difference between the x-Values of the same track at t=0 and t=7 h on unprojected and unsmoothed 3D-data. It was used as a measuring tool instead of speed, as fluctuations in manual tracking strongly affects velocity especially in slow cells.

### Neighbor permanency

is defined as 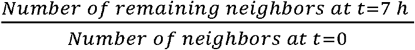. Neighbors are defined as the 6 closest cells to a given cell at t=0. A value of 1 means, that all neighbors were kept.

### Smooth data

As the manual tracking process is fluctuation-prone, we developed a process, taking this uncertainty into account. The *smoothing* process takes every spot as the center of a 10 μm circle and finds the track with the smallest angles, through these areas. The process is reiterated 30 times. Weak phenotypes could potentially lead to false negative results, but false positive phenotypes get much less likely. (Summary and Formula in Fig. S3A, Data after processing: Fig. S3B)

### Mercator projection

[1] An approximation of the central axis is performed by splitting the dataset in 10 subsets along the x-axis. In every subset, yz-coordinates of the center point are approximated by triangulation using the leftmost, rightmost and uppermost points. A central axis is derived from the point of gravity of the first 5 subsets and the last 5 subsets. Based on that, the x-axis is moved with a rotation matrix. This process gets reiterated three times. (Summarized in Fig. S3 B-C)

[2] An yz-vector, 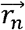 from every point’s respective yz-coordinate to the yz-coordinate of the central axis is generated. Its magnitude is the radius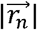 of this point. The maximal radius of all points is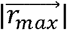. The formula of the central angle *θ* depends on the position of the yz coordinates of every respective point.

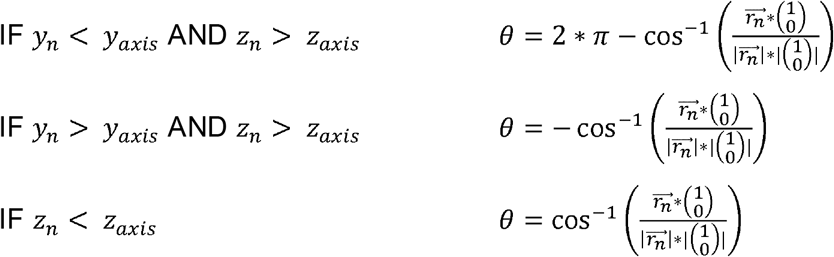

A new y-coordinate is generated using the formula: 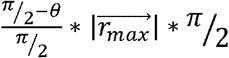 (summarized in Fig. S2 C-E).

[3] To correct the x-axis with respect to 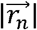, all datapoints are sorted by x-coordinate. 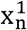 is the x-coordinate of a given spot before correction. 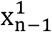 is the point preceeding this point. Its corresponding point after correction is 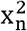 (Summarized in Fig. S3 E-F). For the very first point the formula is: 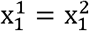

For all further points: 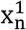: the formula is 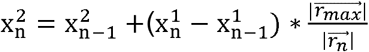

### Track speed mean

was measured in motility lab using smoothed tracking data, in order not to quantify manual tracking inaccuracies.

### Biased angle to x-axis

The usual “biased angle” method measures the bias towards a predefined point. As myotubes do not migrate towards a point, but along a defined axis, we measured the angle-distribution to the x-axis to analyze myotube directionality. As angles get strongly affected by speed, this method can only compare cells with the same “*distance on x”* value (summarized in Fig. S3 F). Rose plots were generated in R using the *ggplot2* package.

### Meandering distance

To compare the directionality of samples with different speeds, their meandering distance 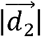 was measured according to the following formula. The median for all tracks on the testis was calculated. 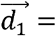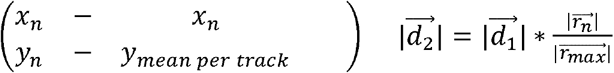

### Cell-matrix adhesion lifetime

was measured with the *spots module* of the Imaris 3.0 software on 2D maximum projections.

### Cell distance over time

between cells isolated by ablation was quantified in R based on Imaris tracking data using the packages *matlib*, *reshape2*, *tibble* and *beeswarm*.

### Statistical Analysis

All statistical tests were performed using Prism 7 (GraphPad). Multiple comparisons were done using parametric or nonparametric Anova, and for single comparisons welsh’s t-test or Mann-Whitney-test was used. Depending on normal distribution, assessed with the Shapiro-Wilk test, either parametric or non-parametric tests were used.

### Image Processing and graphic editing

For image processing and graphic editing, the following software tools were used: Zen Blue (Zeiss), LasX (Leica), Fiji (ImageJ 1.51), Imaris 9.3 (Bitplane), Inkscape 0.91. R Studio 1.2.5042 (RStudio, Inc.) and packages therein mentioned above. For displaying cell tracks, Motility lab was used (Miller, unpublished)

### Computer simulation of testis myoblast behavior

The software was programmed using Unity 2019.2.2f1 (Unity Technologies). A single cell in this model (see also supplementary movie M22) is not simulated as a single agent but consists of multiple simulated protrusion points (black dots). Their geometrical center (centroid) is calculated constantly and constitutes the cells “position”. Protrusion points radially move away from the centroid, mimicking filopodia elongation, but must counter membrane resistance that gets the higher the farther away the point moves from the centroid. On its way every protrusion point creates its own “cell-matrix adhesions” (red dots). They mediate a filopodium (= protrusion points & all its adhesions) static friction which is needed to counter membrane resistance. If membrane resistance is higher than adhesion the entire filopodium gets translated towards the centroid. Protrusion points and cell-matrix adhesions have a lifetime. New protrusion points are generated in a fixed distance from the centroid (grey circle) where the local density is lowest to recapitulate our finding that there is no asymmetry in myotube filopodia assembly. When a protrusion point touches the “adhesion radius” (grey circle) of another cell it loses its cell-matrix adhesions mimicking the measured shortened lifetime of real cell-matrix adhesions. The protrusion point is then turned into an “adhesion point” (green dot) which is recognized by both cells as one of their protrusion points to calculate their respective centroids. For more details see also supplementary material.

## Acknowledgements

We thank the Bloomington Stock Center and VDRC for fly stocks. We thank Thomas Lecuit for providing the Rho1-biosensor. We thank Susanne Önel for providing the Arp3-GFP transgene. We thank Chi-Hon Lee for providing the flies bearing UAS-N-cad-EGFP transgene. We thank Christian Klämbt and Jörg Grosshans for helpful discussion. We thank Caroline Zedler and Franziska Lehne for critical reading of the manuscript. This work was supported by GRK 2213 graduate school program of the Deutsche Forschungsgemeinschaft (DFG) to R. R.-P. and S.B.

## Supplementary material Supplementary figures

**Figure S1.**
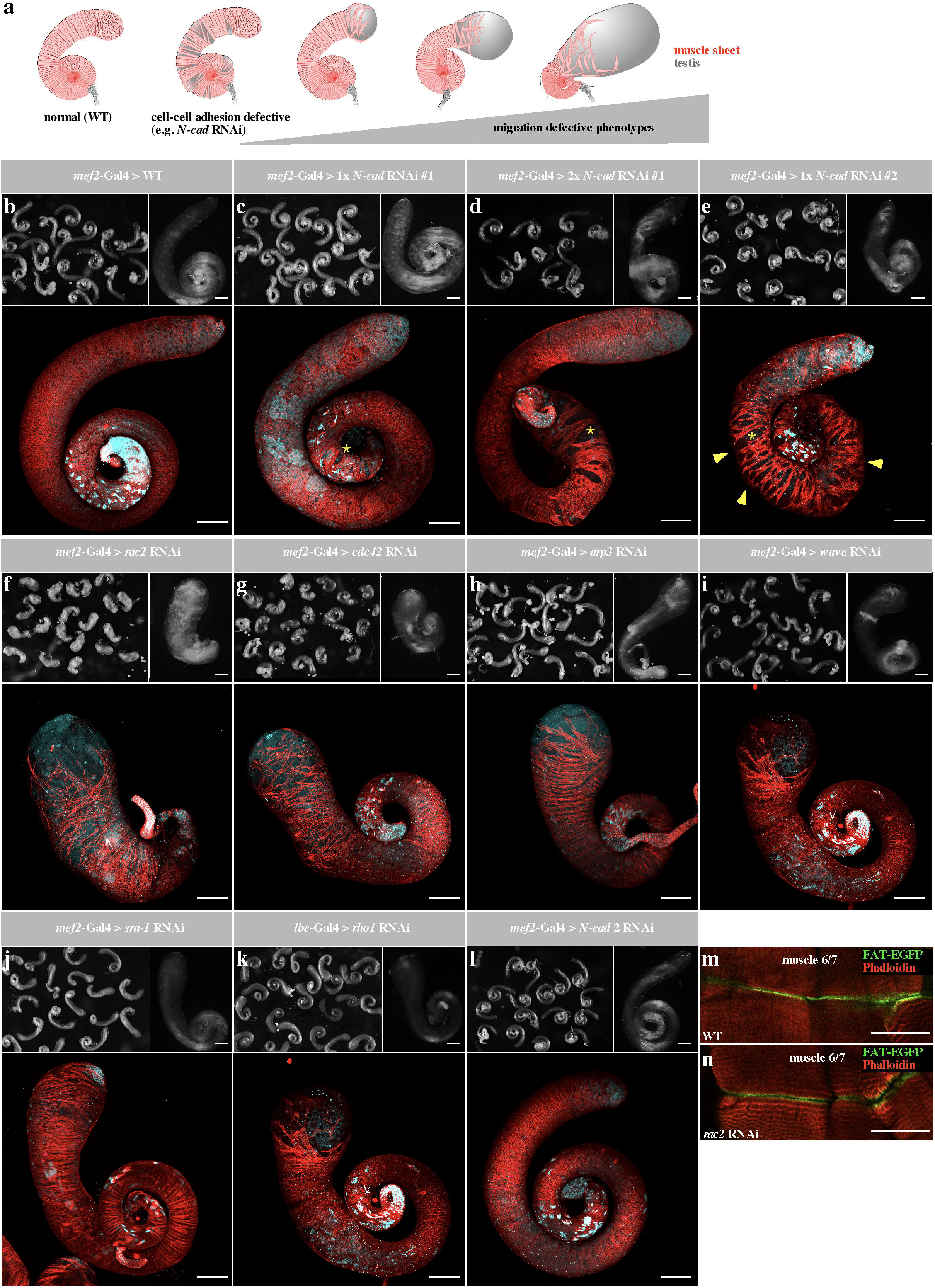
**a.** Graphical representation of types of adult testis defects as a consequence of partial loss of adhesion or migration. **b-l.** Adult testes with different genetic backgrounds. Upper left corner: light micrograph of several testis showing the phenotypic range, upper right corner: light micrograph of a single testis. Bottom: confocal image of a testis. muscle sheet stained with phalloidin (red) and nuclei stained with DAPI (cyan). **b.** Wild type adult testis with a curled shape and an organized and entirely closed muscle sheet. **c, d.** Expression of a *N-cad* RNAi transgene #1 driven by *mef2-Gal4* leads to small holes in the muscle sheet in a dose dependent-manner^30^. **c.** one copy of the RNAi transgene #1; d. two copies of the same RNAi transgene #1. **e.** Expression of a stronger *N-cad* RNAi transgene #2 causes much stronger defects with large holes within the muscle sheet (yellow arrowheads). **f.** *rac2* RNAi driven by *mef2-Gal4* leads to strong migration defects with a strongly dilated tip, partially uncovered, partially covered in disorganized muscles. **g.** *cdc42* RNAi driven by *mef2-Gal4,* resembles *rac2* RNAi with slightly milder defects. **h, i, j.** *arp3*, *wave* and *sra-1* RNAi driven by *mef2-Gal4,* leads to mild migration defects with a slightly dilated tip and small uncovered areas. **k.** *rho1* RNAi driven by *lbe*-Gal4. Even using a weak driver line, prominent migration defects can be observed. **l.** *Ncad2* RNAi driven by *mef2-Gal4* resembles wild type testis without any defects. **m, n.** Confocal images of FAT-GFP driven by *mef2*-Gal4 in larval body wall muscles 6/7. Muscle attachment sites, marked by FAT-GFP are not affected by *rac2* RNAi, indicating that *rac2* depletion has no general impact on matrix adhesion or integrin expression, but specifically affects cell-matrix adhesions in migratory cells. Using sibling flies, no cell-matrix adhesions can be detected in migrating myoblasts.

**Figure S2.**
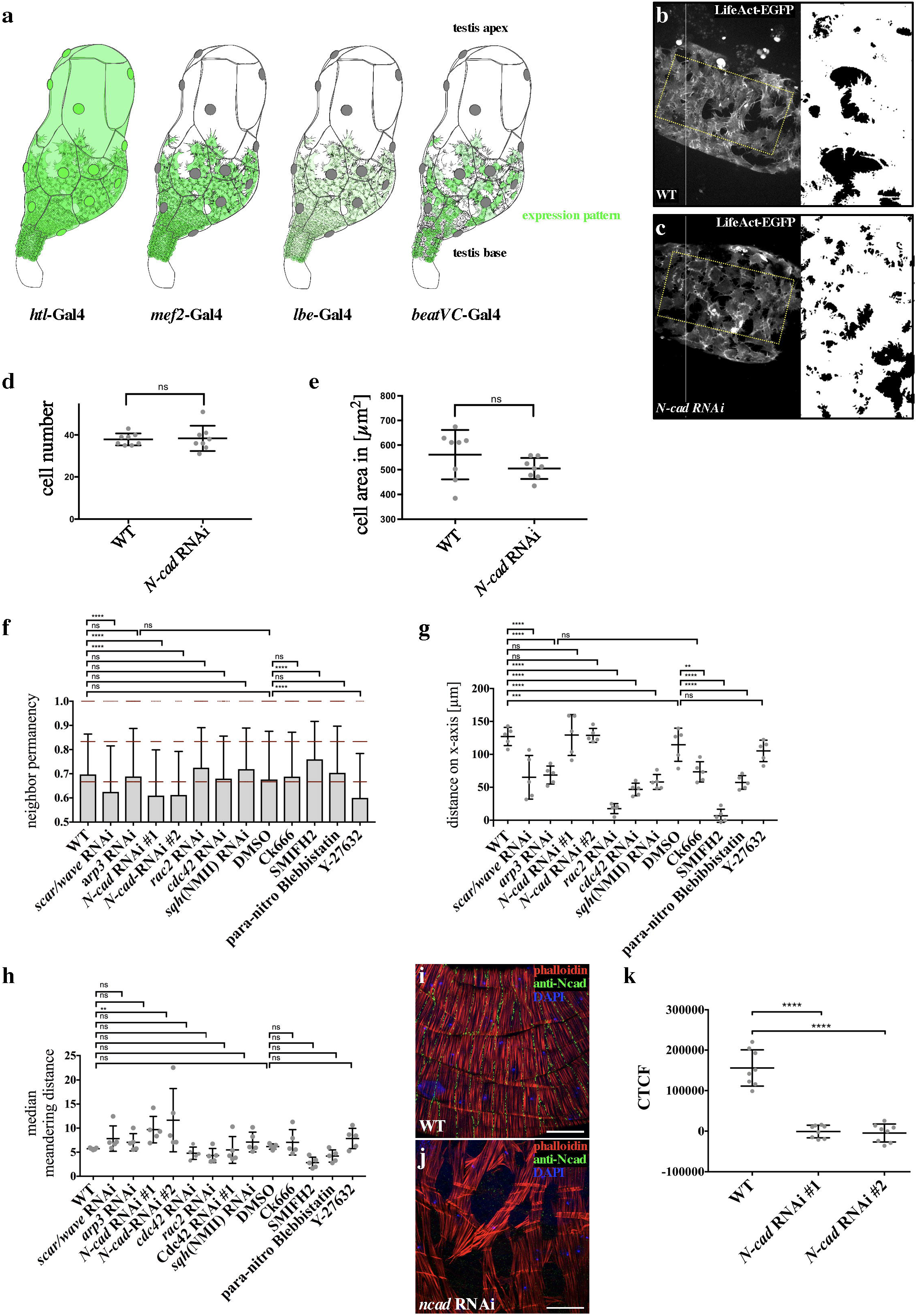
**a.** Graphical representation of the expression patterns of the Gal4 driver lines used (in green). Compare to Fig. 1B. **b, c.** Overview and ROI’s (120 × 220 px) in WT (B) and N-cad RNAi (C) on which quantification in D/E and Fig. 3E, F is based. Marked with yellow dashed lines in the overview**. d.** Cell number inside ROI’s. Cell number is not affected upon *N-cad* RNAi. Source data are provided as a Source Data file. **e.** Area per cell inside ROI’s. Cell Area is not affected upon *N-cad* RNAi. Source data are provided as a Source Data file. **f, g, h, i.** Comparison of neighbor permanency (f.), distance on x-axis (g.), directionality based on biased angle (H) and based on meandering distance (I), for all genotypes. For every genotype, all trackable cells on 5 testes were analyzed. The number of tracks for every genotype equals the number of data points in h. Source data are provided as a Source Data file. **i, j.** Confocal images of adult testis muscle sheet stained with a specific anti-NCad antibody (green), phalloidin (red) and DAPI (blue). **i.** In wildtype (WT) Ncadherin localizes along the cell-cell junction. **j.** Expression of a *ncad* RNAi transgene strongly reduce anti-NCad immunostaining as quantified in **k.** for two independent RNAi transgenes. Source data are provided as a Source Data file.

**Figure S3.**
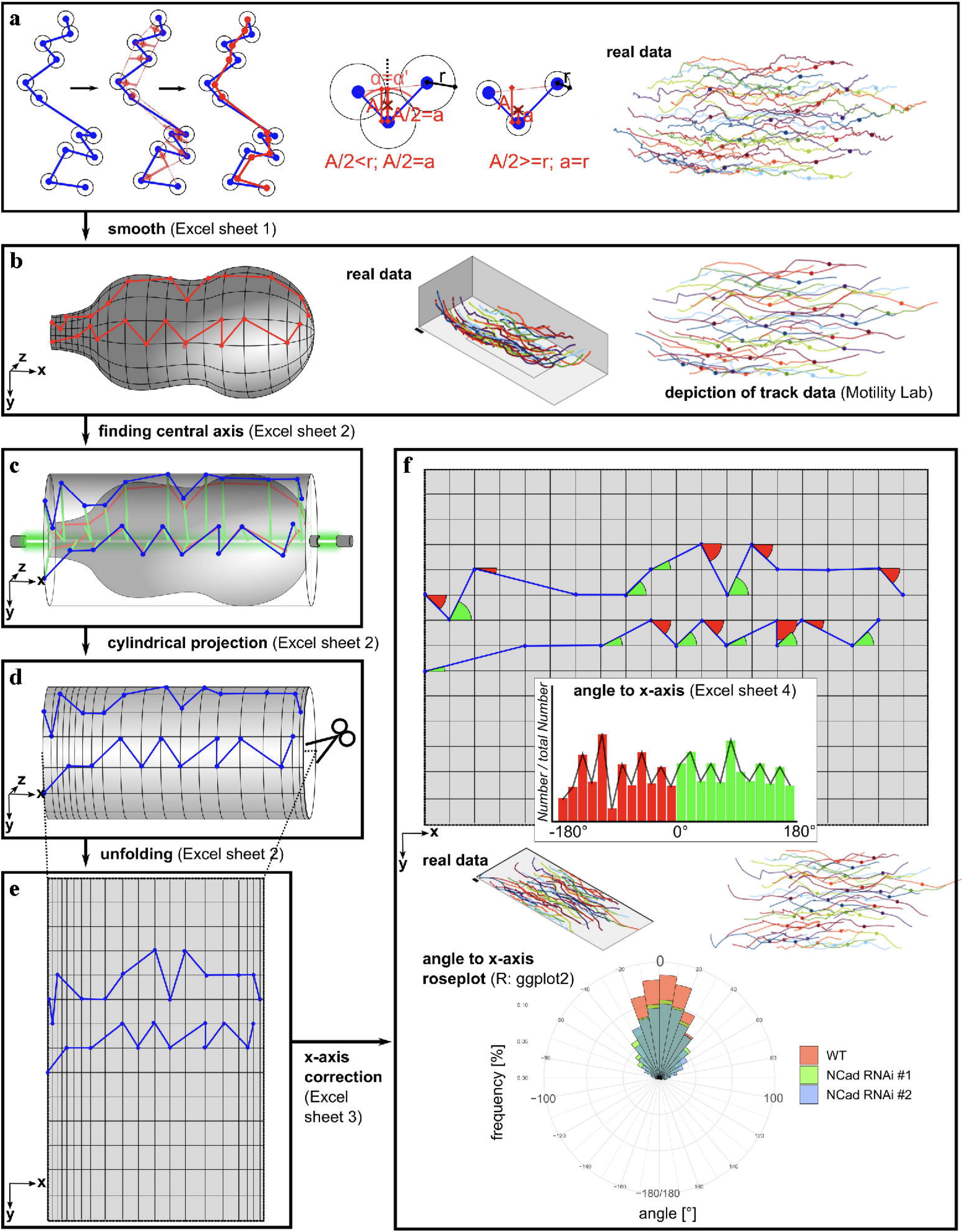
Myotubes migrate on the surface of an ellipsoid, thus on a two-dimensional surface, that is curved in space. This curvature did not allow to apply mathematical rules based in flat geometry. 3D migration tools do not consider the limitations of the surface, to which myotubes are bound but assume they can move freely. Instead, we developed a Mercator projection-based process, which allows for high angle-accuracy but neglects distances. **a.-f.** Steps of Mercator projection are shown (see material and methods for details). Source data are provided as a Source Data file.

**Supplementary table 1.**
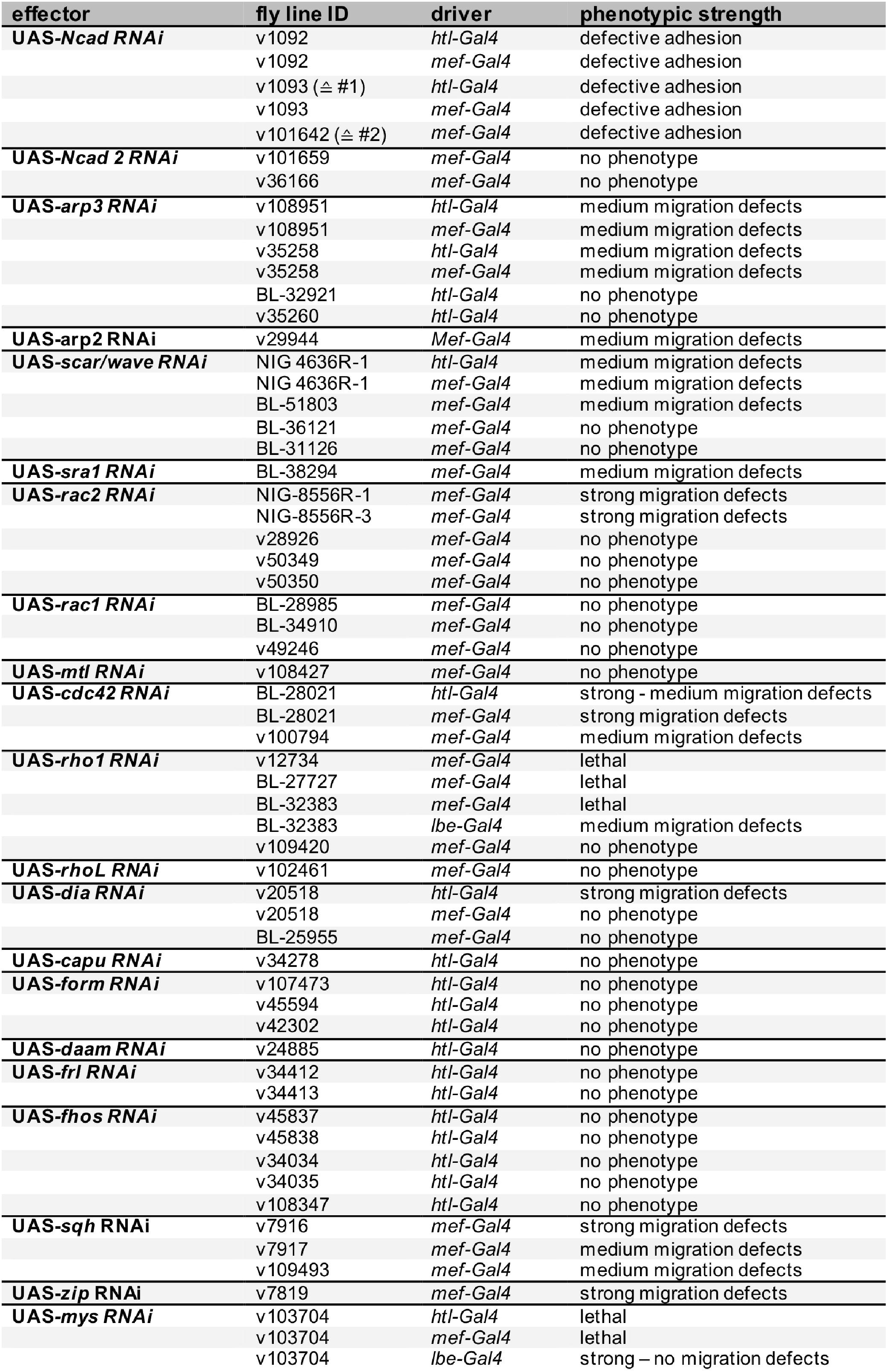
A list of the RNAi transgenes used in this study, and phenotypic strength using different Gal4 driver lines.

## Supplementary methods (simulation model)

Description of the simulation model and details on mathematical modeling.

## Supplementary movies

### Supplementary movie M1

Spinning disc microscopy time-lapse movie of *ex vivo* cultured wild type testes (33h APF) expressing (left) LifeAct-EGFP in myotubes and pigment cells using the htl*-* Gal4 driver, (middle) LifeAct-EGFP only in myotubes using the mef2*-*Gal4 driver, and (right) LifeAct-RFP and a nuclear EGFP in myotubes and pigment cells using the htl*-* Gal4 driver. Scale bar: 50μm.

### Supplementary movie M2

Spinning disc microscopy time-lapse movie of an *ex vivo* cultured wild type testis (33h APF) expressing a LifeAct-EGFP transgene using the *mef2-Gal4* driver. The migration of myotubes was tracked using the Imaris software. An overlay of microscopic data and track data are shown. Scale bar: 30μm.

### Supplementary movie M3

Spinning disc microscopy time-lapse movie of migrating myotubes (marked by LifeAct-EGFP expression) at the front edge of the migrating sheet 60 min in *ex vivo* culture. Scale bar 10 μm.

### Supplementary movie M4

Spinning disc microscopy time-lapse movie of migrating myotubes expressing LifeAct-EGFP and a nuclear EGFP in a mosaic fashion 60 min in *ex vivo* culture, allowing for the analysis of single cells within the migrating sheet. Scale bar 30 μm.

### Supplementary movie M5

Spinning disc microscopy time-lapse movie of an *ex vivo* cultured testis (33h APF) expressing a LifeAct-EGFP transgene, treated with 100 μM CK666. Upon Arp2/3 complex activity inhibition, migration is reduced. Especially cells at the testis base appear to be affected. Scale bar: 50 μm.

### Supplementary movie M6

Spinning disc microscopy time-lapse movie of an *ex vivo* cultured testis (33h APF) co-expressing an *arp3* RNAi together with a LifeAct-EGFP transgene in all myotubes using the *mef2-Gal4* driver. Migration is also mildly reduced by *arp3* RNAi. Scale bar: 50μm.

### Supplementary movie M7

Spinning disc microscopy time-lapse movie of an *ex vivo* cultured testis (33h APF) expressing a LifeAct-EGFP transgene, treated with 10 μM SMIFH2. Myotube migration is completely suppressed. Scale bar: 50μm.

### Supplementary movie M8

Spinning disc microscopy time-lapse movie of migrating myotubes expressing the cell-matrix adhesion reporter FAT-EGFP and a membrane marker Myr-RFP. Scale bar: 10 μm.

### Supplementary movie M9

Spinning disc microscopy time-lapse movie of migrating myotubes expressing the cell-matrix adhesion reporter FAT-EGFP tracked using the Imaris software. Quantified cell-matrix adhesions at the free edge are depicted in red and at the cell-cell edge in green. Scale bar: 20 μm.

### Supplementary movie M10

Spinning disc microscopy time-lapse movie of migrating myotubes expressing the cell-matrix adhesion reporter FAT-EGFP in the middle of the migrating sheet 33 h APF in *ex vivo* culture before and after laser ablation (position is marked by an asterisk). Scale bar: 10 μm.

### Supplementary movie M11

Spinning disc microscopy time-lapse movie of migrating myotubes co-expressing NCad-EGFP and LifeAct-RFP in the middle of the migrating sheet 33 h APF in *ex vivo* culture. Scale bar: 10 μm.

### Supplementary movie M12

Spinning disc microscopy time-lapse movie of *ex vivo* cultured (left) wild type testes compared to (right) testis expressing an *N-cad* RNAi transgene. Myotubes are marked by LifeAct-EGFP transgene expression in all myotubes using the *mef2-Gal4* driver. Scale bar: 50μm.

### Supplementary movie M13

Spinning disc microscopy time-lapse movies of migrating wildtype myotubes (left) compared to myotubes depleted of N-cadherin (right) visualized by the cell-matrix adhesion reporter FAT-EGFP. Scale bar: 10 μm.

### Supplementary movie M13

Spinning disc microscopy time-lapse movie of an *ex vivo* cultured testis (33h APF) co-expressing a *rac2* RNAi together with a LifeAct-EGFP transgene in all myotubes using the *mef2-Gal4* driver. Myotube migration is almost completely disrupted. Scale bar: 50μm.

### Supplementary movie M14

Spinning disc microscopy time-lapse movie of *ex vivo* cultured wild type testis (33h APF) expressing LifeAct-EGFP using the htl*-*Gal4 driver. The isolated single myotube by laser ablation is marked by an asterisk. Scale bar 100 μm.

### Supplementary movie M15

Spinning disc microscopy time-lapse movie of *ex vivo* cultured wild type testis (33h APF) expressing LifeAct-EGFP using the htl*-*Gal4 driver. The isolated myotube pair by laser ablation is marked by an asterisk. Scale bar 100 μm.

### Supplementary movie M16

Spinning disc microscopy time-lapse movie of an *ex vivo* cultured testis (33h APF) co-expressing a *rac2* RNAi together with a LifeAct-EGFP transgene in all myotubes using the *mef2-Gal4* driver. Myotube migration is almost completely disrupted. Scale bar: 50μm.

### Supplementary movie M17

Spinning disc microscopy time-lapse movie of an *ex vivo* cultured (left) wild type testis compared to (right) testis expressing a *cdc42* RNAi transgene. Migration is strongly affected by *cdc42* RNAi. Cells change their shape, generating massive filopodia-like structures, in comparison to WT Scale bar: 50μm.

### Supplementary movie M18

Spinning disc microscopy time-lapse movies of (right) wild type myotubes, (middle) rac2 depleted myotubes and (right) cdc42 depleted myotubes together with a LifeAct-EGFP transgene, 40 min in *ex vivo* culture. Scale bar 20 μm.

### Supplementary movie M19

Spinning disc microscopy time-lapse movies of (right) wild type myotubes, (middle) rac2 depleted myotubes and (right) cdc42 depleted myotubes marked by the cell-matrix adhesion reporter FAT-EGFP, 40 min in *ex vivo* culture. Scale bar 20 μm.

### Supplementary movie M20

Spinning disc microscopy time-lapse movie of migrating myotubes co-expressing the Rho1activity reporter and LifeAct-RFP to visualize protrusion dynamics. Rho-Sensor activity is found in retracting free edge filopodia (yellow arrowheads). Note that the ubiquitously expressed Rho1 sensor also marks ring canals in the *Drosophila* germline cyst (white arrowheads). Scale bar: 10 μm.

### Supplementary movie M21

Spinning disc microscopy time-lapse movies of *ex vivo* cultured (from left to right) testis treated with (A) the Rok inhibitor (Y27632), (B) treated with the blebbistatin, expressing a (C) *sqh* and (D) *zip* RNAi transgene. Migration is strongly affected. Scale bar: 50μm.

### Supplementary movie M22

Computer simulation model. Cells were positioned at one end of a confinement roughly mimicking the unfolded testis surface. **(A)** If cells behave according to the default setting, meaning that just adhesion but not filopodia lifetime is affected by contact, then all free space gets covered while cells keep their cohesion. **(B)** Isolated cells keep their cohesion as well, but randomly migrate, until they are contacted by the expanding sheet. After contact, they move along, as part of the sheet. **(C)** When filopodia disassemble shortly after contact, the behaviour resembles CIL, as there is a short phase of protrusion asymmetry, shifting the centroids apart from each other. Shortly after contact the repulsive motion ceases, as new filopodia emerge, until another cell moves close. All space gets covered, but there is no cohesion between cells. (A-C) In all scenarios a clumping of cells can be observed at the testis base. It seems to be a consequence of the interaction with the barrier which has no counterpart in a three dimensional limitless but finite surface. It is thereby an artefact with no meaning for the simulation (for more details about the simulation see supplementary material).

